# Allosteric Inhibition of the T Cell Receptor by a Designed Membrane Ligand

**DOI:** 10.1101/2022.08.19.503518

**Authors:** Yujie Ye, Shumpei Morita, Kiera B. Wilhelm, Jay T. Groves, Francisco N. Barrera

## Abstract

The T cell receptor (TCR) is a complex molecular machine that directs the activation of T cells, allowing the immune system to fight pathogens and cancer cells. Despite decades of investigation, the molecular mechanism of TCR activation is still controversial. One of the leading activation hypotheses is the allosteric model. This model posits that binding of pMHC at the extracellular domain triggers a dynamic change in the transmembrane (TM) domain of the TCR subunits, which leads to signaling at the cytoplasmic side. We sought to test this hypothesis by creating a TM ligand for TCR. Previously we described a method to create a soluble peptide capable of inserting into membranes and bind to the TM domain of the receptor tyrosine kinase EphA2 (Alves *et al., eLife* 2018). Here we show that the approach is generalizable to complex membrane receptors, by designing a membrane ligand for TCR. We observed that the designed peptide caused a reduction of Lck phosphorylation of TCR at the CD3ζ subunit. As a result, in the presence of this Peptide Inhibitor of TCR (PITCR), the proximal signaling cascade downstream of TCR activation was significantly dampened in T cells. Co-localization and co-immunoprecipitation results in DIBMA native nanodiscs confirmed that PITCR was able to bind to the TCR. We propose that PITCR binds into a crevice present between the TM helices of the CD3ζ and CD3ε(δ) subunits. Our results additionally indicate that PITCR disrupts the allosteric changes in the compactness of the TM bundle that occur upon TCR activation, lending support to the allosteric TCR activation model. The TCR inhibition achieved by PITCR might be useful to treat inflammatory and autoimmune diseases and to prevent organ transplant rejection, as in these conditions aberrant activation of TCR contributes to disease.

## Introduction

T cells are central players in the adaptive immune response. Different types of T cells recognize the presence of pathogenic organisms and cancer cells and orchestrate diverse immune activities intended to kill the damaging cells (Courtney et al., 2017; Ganti et al., 2020). The T cell receptor (TCR) is a protein complex present at the membrane of T cells that allows to detect the presence of foreign molecules. The TCR engages with antigen-presenting cells (APC), where peptide fragments are displayed at the major histocompatibility complex (pMHC) (Chakraborty & Weiss, 2014). Recognition of pMHC by the TCR triggers an intricate signaling cascade that activates the T cell response (Courtney et al., 2018; Kuhns & Davis, 2008). In αβ T cells, pMHC binding occurs at the TCRαβ subunits. The TCR complex additionally contains four types of CD3 subunits: ε, γ, δ and ζ, forming the TCR-CD3 complex (referred herein as TCR). The dominant stoichiometry of TCR is composed of TCRαβ–CD3εγ–CD3εδ–CD3ζζ (Call et al., 2002; Dong et al., 2019; Mariuzza et al., 2020). The CD3 subunits relay the information of the pMHC binding event across the membrane, initiating TCR proximal signaling. TCR downstream signaling cascade starts by phosphorylation of ITAMs (immunoreceptor tyrosine-based activation motifs) present in all CD3 subunits, particularly at ζ, which leads to a transient increase in calcium levels in the cytoplasm (Au-Yeung et al., 2018; Lo et al., 2018).

Despite decades of effort, there is not yet a clear understanding of how TCR is activated after recognition of pMHC (Mariuzza et al., 2020). A number of different activation modes have been proposed, namely the aggregation/clustering model, the segregation model, the mechanosensing model, and the allosteric model. In this latter functional hypothesis, the signal resulting from binding of pMHC to the extracellular region of the TCRαβ is dynamically transmitted across the transmembrane (TM) region into the ITAMs. While there is growing evidence supporting the allosteric model (Chen et al., 2022; Schamel et al., 2019), the molecular mechanism of the allosteric triggering of TCR-CD3 is not clear. Recent reports provide a plausible mechanistic framework on how allosteric changes affect the TM helical bundle: TCR activation involves a quaternary relaxation of the TM helical bundle, whereby in the active state there is a loosening in the interaction between the TM helices of the TCRβ and CDζ (Lanz et al., 2021; Prakaash et al., 2021).

Here, we have used rational design to develop a peptide (PITCR) to target the TM region of TCR. PITCR comprises the TM domain of the ζ subunit (Call et al., 2006) modified by the addition of acidic residues to convert it to a conditional TM sequence. The PITCR peptide is used to test the allosteric relaxation model, as its binding to the TM region of TCR can be reasonably expected to alter the conformation and/or dynamics/packing of the helical bundle. We observed that PITCR robustly inhibited the activation of the TCR. The results obtained in this work support the allosteric relaxation activation model and provide new mechanistic insights into TCR activation.

## Results

### PITCR decreases phosphorylation of the ζ chain upon TCR activation

We recently reported an approach to transform the isolated TM domains of human receptors into peptides that function as conditional TM sequences: they are highly soluble in water, while they have the ability to insert into the membrane in the TM orientation that allows the peptide to interact laterally with their natural binding partners (Alves et al., 2018). We applied this protocol to the CD3ζ TM, to generate the PITCR peptide. Biophysical experiments in synthetic lipid vesicles showed that the design for PITCR was successful, as it was soluble in aqueous solution and able to insert into membranes (Figure 1-Figure supplementary 1).

The TCR at the surface of human Jurkat T cells is activated upon binding of the monoclonal antibody (mAb) OKT3, which has been widely applied to study T cell signaling (Lo et al., 2018; Lo et al., 2019). TCR activation is initiated by phosphorylation of tyrosine residues at the ITAMs of the ζ chain by Lck (lymphocyte-specific protein tyrosine kinase) (Figure 1A) (Courtney et al., 2017). To investigate whether PITCR affects TCR activation, we treated Jurkat cells with PITCR before stimulation with OKT3. The immunoblot results revealed that PITCR reduced phosphorylation of the ζ chain at residues Y142 and Y83 after TCR activation (**Figure 1B-C and Figure 1 - Figure supplementary 2**). These results suggest that PITCR binding reduces activation of the TCR.

**Figure 1.**
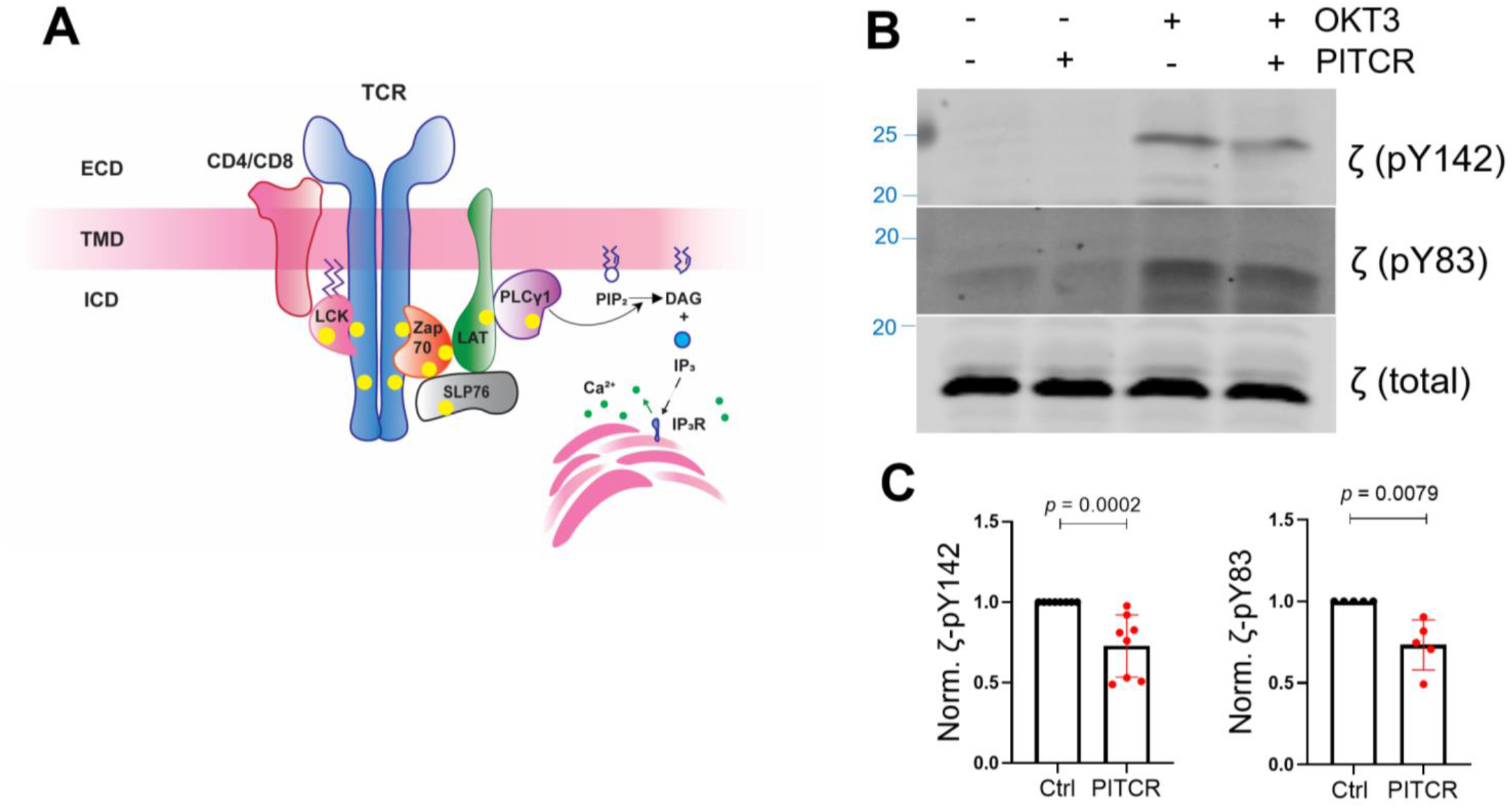
PITCR reduces phosphorylation of the *ζ* chain in response to OKT3. **(A)** Cartoon that illustrates TCR proximal downstream signaling. The plasma membrane is shown as a horizontal bar, and phosphorylation sites are shown as yellow dots. ECD: extracellular domain; TMD: transmembrane domain; ICD: intracellular domain of TCR. **(B)** Jurkat cells were treated with PITCR, followed by stimulation with OKT3. Lysates were analyzed by immunoblot to detect TCR phosphorylation of ζ (pY142 and pY83). Total ζ levels were assessed and no change was observed. Data are representative at least five independent experiments. **(C)** Quantification of phosphorylation at both tyrosine residues in the presence of OKT3, normalized to data in the absence of PITCR (based on data from Figure 1 - Figure supplementary 2). Error bars are the SD. *p* values were calculated using a two-tailed Mann Whitney test.

### Phosphorylation of TCR proximal signaling molecules is downregulated by PITCR

Since PITCR inhibited TCR phosphorylation after activation, we sought to explore the effect of PITCR on TCR downstream signaling. TCR activation induces the recruitment of Zap70 (ζ chain associated protein kinase 70) to the phosphorylated TCR, where it is itself phosphorylated by Lck **(Figure 1A)**. The activated Zap70 then phosphorylates LAT (linker for activation of T cells) and SLP76 (SH2 domain containing leukocyte protein of 76 kDa), and as a result PLCγ1 (phospholipase Cγ1) is recruited and phosphorylated (Courtney et al., 2018; Lo et al., 2018; Lo & Weiss, 2021). In agreement with the hypothesis that PITCR inhibits TCR activation, in the presence of peptide we observed a statistically significant decrease in the phosphorylation of Zap70, LAT, SLP76 and PLCγ1 (**Figure 2 and Figure 2 - Figure supplementary 1**). However, PITCR did not affect phosphorylation of Lck (**Figure 2-Figure supplementary 2**). Since basal phosphorylation of Zap70, LAT, SLP76 and PLCγ1 was observed in the absence of OKT3 (**Figure 2A**), we could determine that the effect of PITCR is specific to stimulation of the TCR.

**Figure 2.**
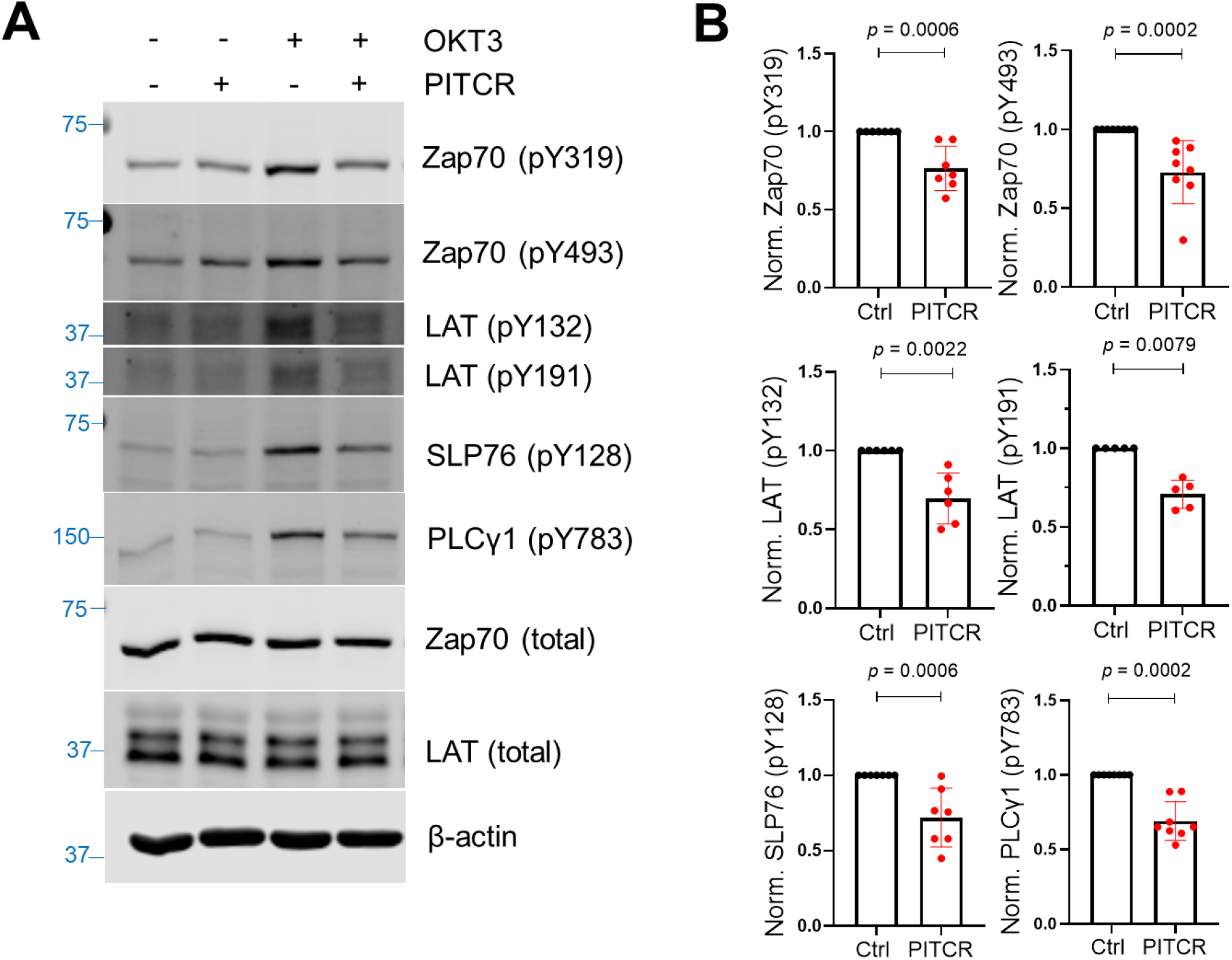
PITCR reduces phosphorylation of TCR proximal signaling proteins after activation. **(A)** Immunoblot analysis of lysates to detect phosphorylation of Zap70 (pY319 and pY493), LAT (pY132 and pY191), SLP76 (pY128), and PLCγ1 (pY783). Total protein levels of Zap70, LAT and the housekeeping protein β-actin were assessed, revealing no changes. Data are representative of at least five independent experiments. **(B)** Quantification of phosphorylation in the presence of OKT3, normalized to data in the absence of PITCR (based on data from Figure 2-Figure supplementary 1). Error bars are the SD. *p* values were calculated using a two-tailed Mann Whitney test.

Our data therefore indicate that PITCR causes a robust inhibition of the proximal signaling cascade triggered by TCR activation.

### PITCR reduces intracellular calcium response

After TCR activation, the active PLCγ1 hydrolyzes phosphatidyl inositol 4,5-bisphosphate (PIP_2_) to generate inositol trisphosphate (IP_3_) and diacylglycerol (DAG). Free IP3 diffuses across the cytoplasm and binds to the IP3 receptor at the endoplasmic reticulum (ER), causing the release of calcium from ER storage (Figure 1A) (Courtney et al., 2018; Lewis, 2001; Trebak & Kinet, 2019). Based on our previous results, we reasoned that PITCR should inhibit the cytoplasmic calcium influx that follows TCR activation. We tested this idea with a kinetic analysis of intracellular calcium release using the calcium indicator Indo-1 (Lo et al., 2018).

Consistent with our expectation, we observed that the calcium response after OKT3 stimulation was attenuated in the PITCR treatment group compared with control conditions (**Figure 3 A and D, Figure 3 - Figure supplementary 1**). We used as a negative control pHLIP (Scott et al., 2019; Scott et al., 2017), a different conditional TM peptide that is not expected to interact with membrane proteins (Alves et al., 2018). The dynamic calcium curve of pHLIP-treated group was within the error of the control curve (**Figure 3 B and D**). To further test specificity of PITCR, we performed a mutation (replacing Gly41 for a Pro) in PITCR that is expected to form a helical kink and disrupt the TM helix. Control biophysical experiments indicated that the G41P mutation did not prevent the peptide to act as a conditional TM (**Figure 1 - Figure supplementary 1**). We observed that PITCRG41P was unable to inhibit the calcium influx in response to OKT3 treatment (**Figure 3 C and D**). These data indicate that PITCR specifically impaired the calcium response that occurs in Jurkat cells after TCR activation, in agreement with the observed decrease in phosphorylation of PLCγ1 (**Figure 2A**).

**Figure 3.**
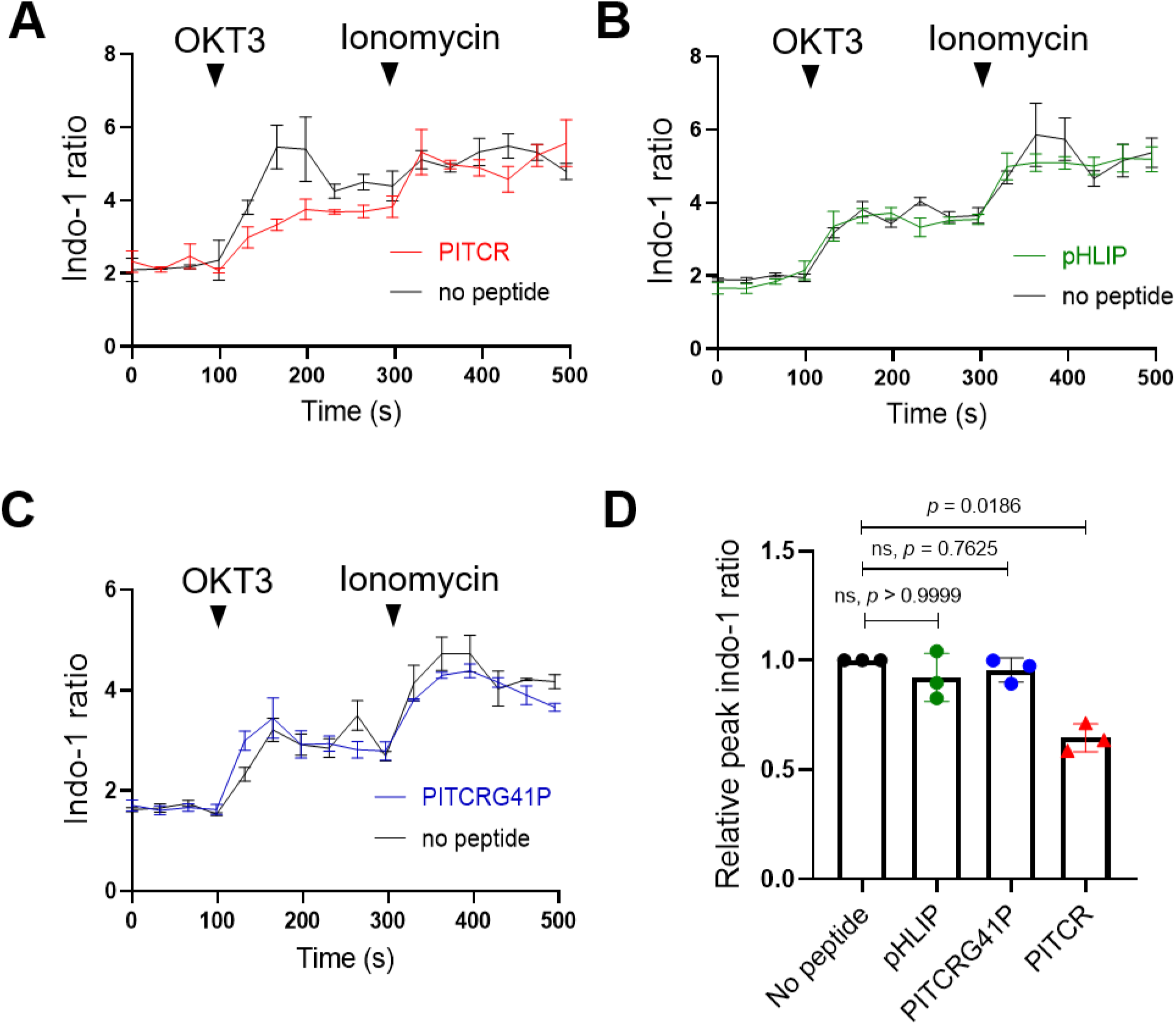
PITCR reduces the TCR intracellular calcium response. Jurkat cells were stained with the fluorescent dye indo-1 AM, followed by treatment with PITCR **(A)**, pHLIP as a negative control **(B)**, or the variant PITCRG41P **(C)** and stimulated with OKT3. Ionomycin was applied as a positive calcium control. The indo-1 ratio was calculated from fluorescence at 405 nm (calcium-bound) divided by 475 nm (calcium-free). Data are representative of three independent experiments. Each independent experiment includes at least four technical replicates. Error bars are the SEM. **(D)** Quantification of the maximum Indo-1 increase after OKT3 activation, normalized to no peptide treatment. Error bars are the SD. *p* values were calculated from a Kruskal-Wallis test with Dunn’s multiple comparisons test.

### Inhibition of TCR activation by antigen presenting cells

While the OKT3 mAb efficiently stimulates TCR signaling, we sought to test the effect of TCR in a more physiologically relevant TCR activation, consisting of T cell stimulation by binding to pMHC in antigen presenting cells (APCs) (Lo et al., 2018). Similar to human primary cytotoxic CD8^+^ T cells, OT1^+^-TCR CD8^+^ Jurkat T cells (J.OT1.CD8) can recognize the ovalbumin (OVA) peptide presented by H-2K^b^-MHC I on T2 APC cells (T2K^b^) (**Figure 4 A and B**). TCR engagement results in increased levels of CD69, a T cell activation marker (Lo et al., 2018; Lo et al., 2019; Wolpert et al., 1997).We treated J.OT1.CD8 cells with PITCR followed by incubation with T2K^b^ cells pre-treated with a range of OVA concentrations and measured the CD69 expression using flow cytometry (**Figure 4 C and D**). As expected, in the presence of high OVA concentrations we observed CD69 upregulation (**Figure 4 C**). Our data showed that PITCR caused a significant reduction of CD69 levels (**Figure 4 C and D**). To further examine the specificity of PITCR, we again used pHLIP as a negative control, and we observed that pHLIP did not elicit significant changes. Our data therefore indicate that PITCR specifically impaired CD69 upregulation in J.OT1.CD8 cells in response to OVA stimulation. These results show that PITCR also achieves robust inhibition when TCR is activated by binding to pMHC in APC.

**Figure 4.**
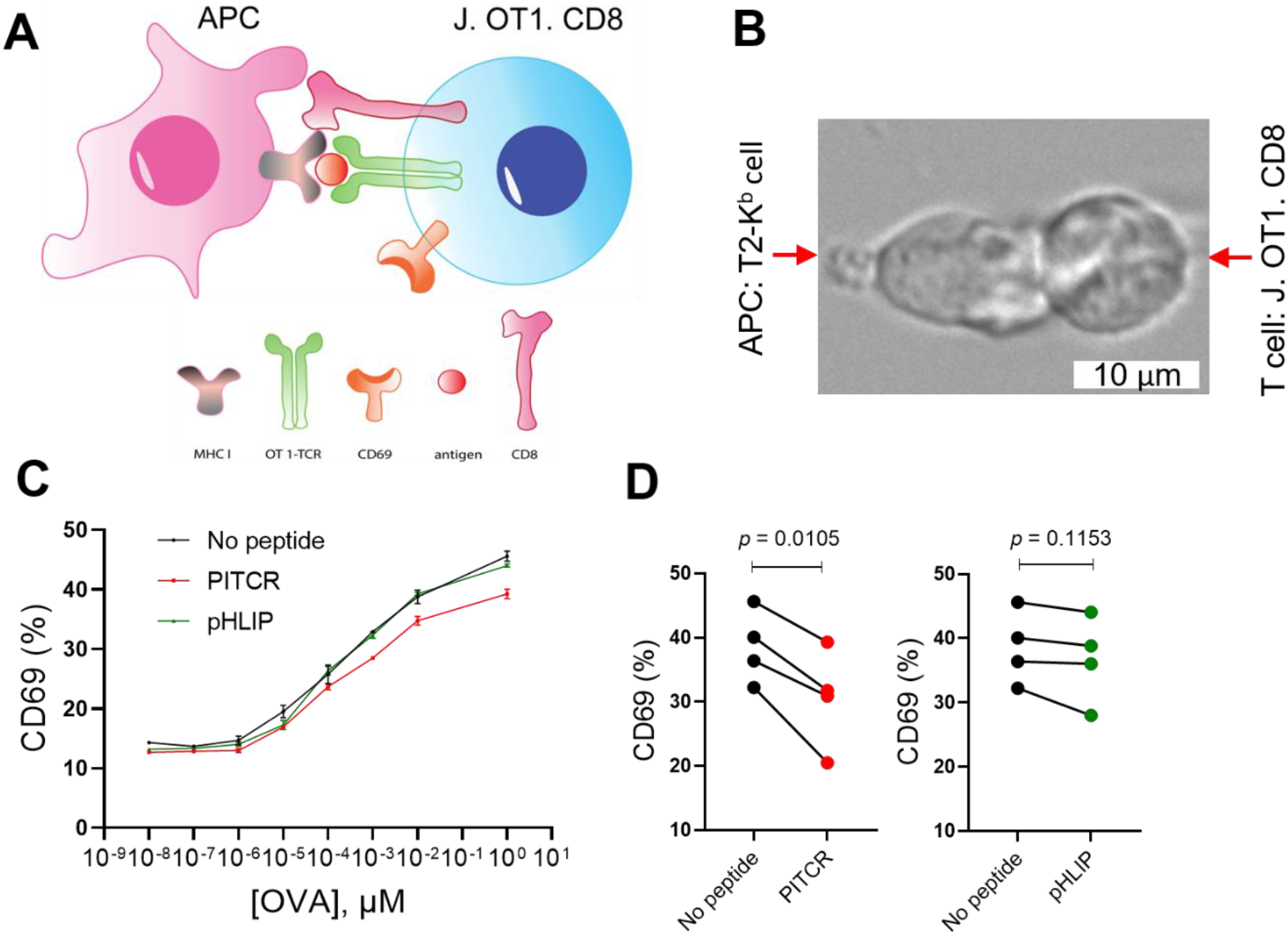
PITCR reduces CD69 expression after T cell activation by APC. **(A)** Cartoon showing T cell interaction with APC. **(B)** A live cell microscopy image that depicts engineered Jurkat OT1^+^ TCR-CD8^+^ T cells interacting with T2K^b^ APC pre-incubated with the peptide antigen OVA. **(C)** Jurkat OT1^+^ TCR-CD8^±^ T cells were treated with PITCR or pHLIP (as a negative control), and then incubated with T2K^b^ cells at different concentrations of OVA, followed by CD69 flow cytometry analysis. The upregulation of CD69 is representative of four independent experiments. Each independent experiment includes two technical replicates. Error bars are the SD. **(D)** Quantification of CD69-positive cells at [OVA] = 1 μM for PITCR (red) and negative control pHLIP (green). Each dot pair represents one independent experiment. *p* values were calculated from two-tailed paired *t*-test.

### PITCR co-localizes with the TCR in Jurkat cells

Our results in Jurkat cells clearly show that PITCR reduces TCR activation. We sought next to determine if this was a specific effect that resulted from a direct interaction between the peptide and TCR. First, we performed co-localization experiments in Jurkat cells. To this end, we fluorescently labeled PITCR with dylight680 (PITCR680). We employed super-resolution confocal microscopy with lightning deconvolution to investigate PITCR680 co-localization with TCR, as detected with an antiCD3 ε antibody. We observed that PITCR680 localized to some areas of the Jurkat cells, corresponding to the plasma membrane and intracellular organelles, probably endosomes (**Figure 5 A**). In these two regions we observed co-localization between PITCR680 (magenta) and TCR (CD3ε, green) (**Figure 5 A**). To better assess co-localization, we plotted graphic profile curves on regions of interest (ROI), which revealed clear overlap in some areas. To quantify the degree of co-localization, we calculated the Mander’s M1 coefficient, which showed a value of ~0.8 (**Figure 5 D**). This result reveals that around 80% of PITCR680 signal overlaps with the TCR. We also observed robust co-localization using the Pearson correlation coefficient (*r* = ~0.4) (**Figure 5 – Figure supplementary 2**) (Costes et al., 2004). To further explore whether TCR activation influences co-localization, we stimulated PITCR680-treated Jurkat cells with OKT3. We observed similar results, suggesting that PITCR is able to bind to TCR before it is activated by OKT3.

**Figure 5.**
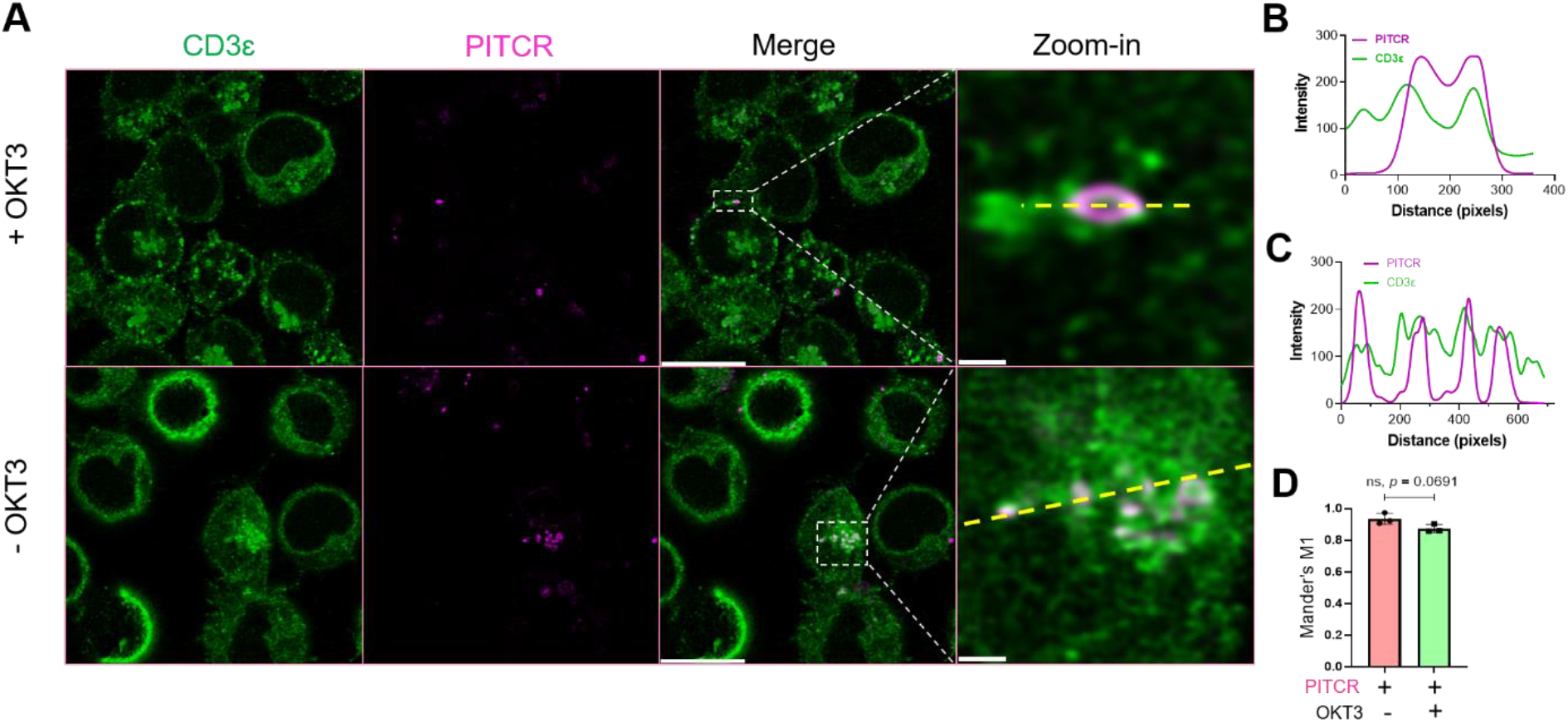
PITCR colocalizes with TCR. **(A)** PITCR680 and CD3ε colocalization was studied in the presence and absence of activation with OKT3. Scale bars = 10 μm. Representative areas with colocalization at the plasma membrane (*top*) and cytoplasm (*bottom*) were zoomed-in, where scale bars are 0.5 μm and 1 μm, respectively. Confocal images are representative of three independent experiments. **(B)** and **(C)** show graphic profile curves (dotted yellow lines) plotted across the zoom-in ROI in + OKT3 and OKT3, respectively. Magenta lines denote PITCR, and green lines denote CD3ε. Overlap indicates colocalization. **(D)** Quantification of colocalization by Mander’s coefficient (M1), corresponding to the fraction of PITCR that overlaps with CD3ε. Error bars indicate SD. *p* value was calculated from two tailed unpaired *t* test.

### Co-localization of PITCR with ligand-bound TCR in primary murine cells

Primary murine CD4^+^ T cells provided an orthogonal method to assess co-localization of the peptide with TCR. Splenocytes from the TCR(AND) mice, hemizygous for H2k, were pulsed with 1 μM moth cytochrome c (MCC) peptide and cultured with the T cells for two days. The T cell blasts were treated with IL-2 from the day after harvest to the fifth day after harvest, at which point the cells were used in experiments. T cells in this state respond to antigen with single-molecule sensitivity. T cells treated with either PITCR or a vehicle control were stimulated by contact with supported bilayers functionalized with agonist pMHC (MCC peptide labeled with Atto647N) and the adhesion molecule ICAM-1 (Lin et al., 2019) (McAffee et al., 2021). The primary mouse T cells activated normally upon treatment with PITCR as measured by NFAT translocation (**Figure 6-Figure supplementary 1**) (Lin et al., 2019). We performed surface-selective imaging by total internal reflection fluorescence microscopy and immune synapse formation was imaged (Biswas & Groves, 2019; Grakoui et al., 1999; Mossman et al., 2005; Yu et al., 2012).TCR-pMHC complexes were selectively distinguished from free pMHC using an elongated image exposure time strategy (Lin et al., 2019; O’Donoghue et al., 2013; Pielak et al., 2017). We observed the c-SMAC (central supramolecular activation cluster) structure (**Figure 6**), as previously reported for activated T cells (Bromley et al., 2001; Grakoui et al., 1999). PITCR conjugated to AZDye 555 (PITCR555) could be detected in intracellular structures (**Video 1**), consistent with confocal microscopy results. Additionally, a distinct population of plasma-membrane-bound peptide could be also observed in some cells (**Figure 6**). In these cases, PITCR exhibited c-SMAC localization together with TCR-pMHC complexes. Although the image resolution was insufficient to definitively confirm molecular binding between PITCR and TCR-pMHC complex, their co-localization is suggestive of interaction. We note that PITCR is a partial TCR inhibitor, and thus is not expected to significantly block c-SMAC formation or activation in these primed mouse T cells due to their extreme sensitivity to antigen. Additionally, TM-TM interactions are often highly sensitive to mutations (He et al., 2017; Westerfield et al., 2021). Several TM residues in the CD3 subunits that according to our model interact with TCR are different in the murine and the human amino acid sequences. Therefore, we expect PITCR to be less efficient targeting the mouse TCR. In aggregate, the co-localization experiments support binding of PITCR to TCR in Jurkat and primary T cells.

**Figure 6:**
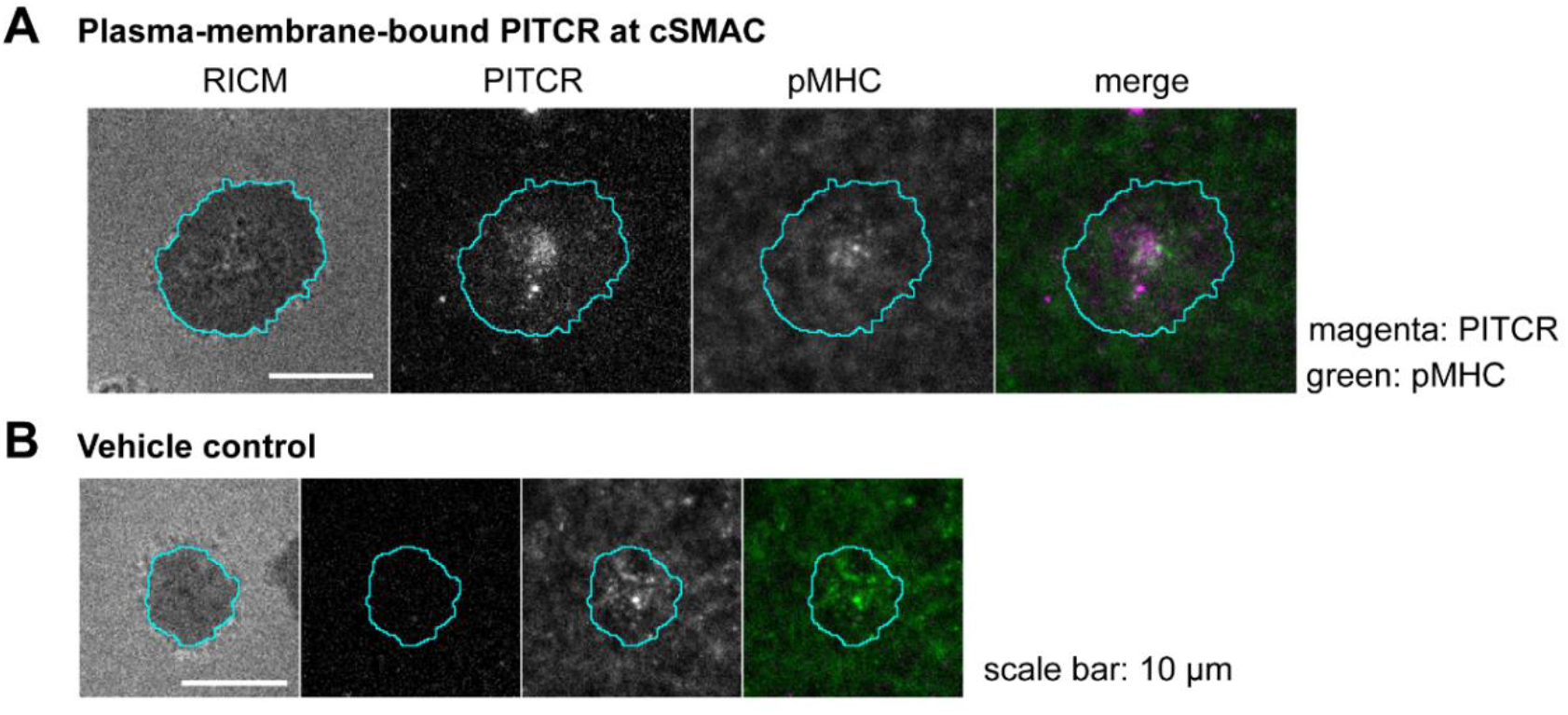
Colocalization of PITCR and the TCR-pMHC complex in primary murine CD4^+^T cells. **(A)** Images of plasma-membrane-localized PITCR555 and TCR-pMHC complex in a representative T cell adhering to supported lipid bilayer functionalized with pMHC (19-23 molecules/μm^2^, 9% labeled with Atto647N) and ICAM-1 (~20 molecules/μm^2^). TCR-pMHC complex was selectively visualized with a long exposure time (500 ms). PITCR exhibited localization at c-SMAC together with the TCR-pMHC complex. **(B)** Vehicle control showed no signal at the PITCR channel.

### PITCR interacts with the TCR in Jurkat cells

We sought to determine if the observed co-localization indeed resulted from binding between PITCR and TCR. We developed an assay to maintain the TCR complex of Jurkat cells in a native lipid environment, consisting of using the polymer diisobutylene maleic acid (DIBMA) to form native nanodiscs. On these samples we performed a co-immunoprecipitation (Co-IP) experiment using an anti-CD3 ε antibody (UCHT1). We observed bands of TCR β, CD3 ε and CD3 ζ in the anti-CD3 ε Co-IP lysates (**Figure 7**), indicating that TCR had been successfully immunoprecipitated after solubilization with DIBMA. When cells were incubated with PITCR680, we observed in the Co-IP samples a fluorescent band of molecular weight (~5 kDa) similar to that of PITCR680 (4.8 kDa) (**Figure 7**). To further explore whether TCR activation would affect binding of PITCR680 to the complex, we applied OKT3 as previously described. We observed the ~5 kDa fluorescent band as well. These results indicate that PITCR680 interacts with the TCR irrespective of activation by OKT3, in agreement with the co-localization results in Jurkat cells (**Figure 5**).

**Figure 7.**
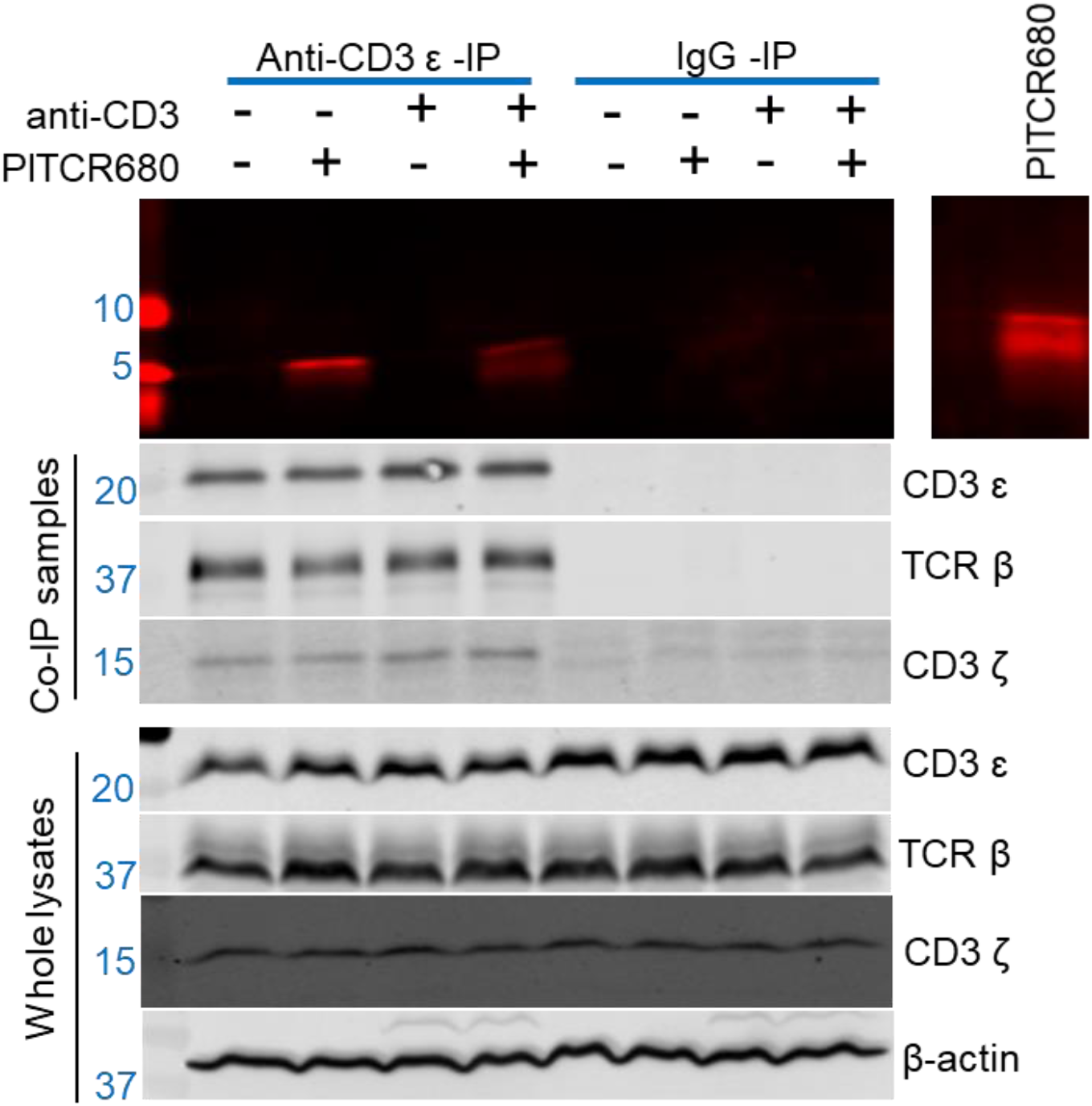
PITCR interacts with TCR. Jurkat cells were treated with PITCR-680. TCR nanodiscs were immunoprecipitated with the mAb UCHT1, or with IgG as a negative control. Fluorescent detection of PITCR680 after CoIP or run in the gel directly as a positive control (*right side panel*). Below are shown immunoblot analysis of Co-IP samples and whole lysates to probe CD3 ε, TCR β and CD3 ζ. β actin was a loading control. Data are representative of at least three independent experiments.

### PITCR weakens the interaction of the ζ subunit with the rest of the complex after TCR activation

We next sought to understand the mechanism by which PITCR partially inhibits TCR activation. It has been recently proposed that TCR activation involves a change in robustness of the TM helical bundle. This allosteric change can be detected by immunoprecipitation after solubilization in DDM, as a loose complex is less resistant to this detergent (Lanz et al., 2021; Prakaash et al., 2021). We optimized this assay for our experimental conditions and investigated the interaction between the ζ chain and rest of the TCR complex. We first examined the immunoprecipitated levels of CD3 ε, CD3 ζ and TCR β in the presence of DDM to evaluate the specificity and efficacy of the use of the anti-CD3 ε antibody for our immunoprecipitatation assay. We observed the targeted bands in the immunoblot results for anti-CD3 ε-IP, and no bands in the negative control IgG-IP (**Figure 8A**). These results suggest that our protocol successfully IPs the TCR complex. Once we validated our method, we tested the effect of OKT3 stimulation and PITCR treatment. We observed that OKT3 increased the levels of CD3ζ when compared to both TCRβ2 and CD3ε (ζ */* β2 and ζ */* ε) (**Figure 8**). These results suggested a more robust ζζ interaction with the rest of complex in response to OKT3 stimulation. Based on this result, we reasoned that PITCR could act by reversing the changes in quaternary robustness that occur in the membrane region upon TCR activation. In agreement with our hypothesis, we observed that the ζ */* β2 ratio was decreased when cells were incubated with PITCR and OKT3. However, ζ / ε did not change (**Figure 8**). Taken together, these results suggested that PITCR disrupts the allosteric changes in TM quaternary robustness that occur upon TCR activation.

**Figure 8.**
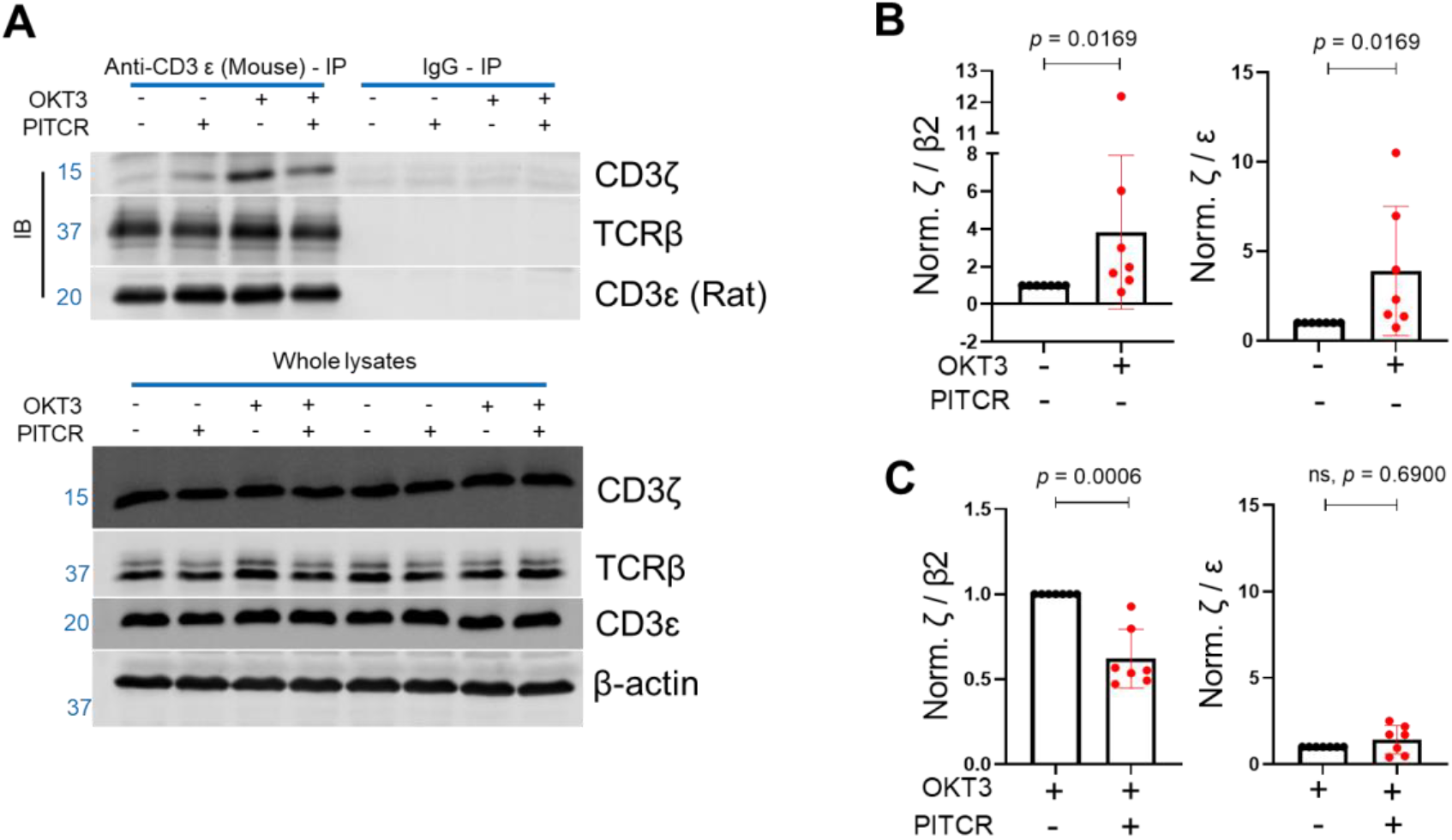
PITCR weakens the interaction of the ζ chain with the rest of the complex after TCR activation. **(A)** Immunoblot analysis of immunoprecipitated samples and whole lysate samples solubilized with DDM. Data are representative of at least three independent experiments. **(B)** Quantification of ζ / β2 and ζ / ε after OKT3 stimulation. **(C)** ζ / β2 and ζ / ε in PITCR-treated OKT3 samples, normalized to OKT3 stimulation. The β2 subunit was studied since it is the most abundant β subunit at the plasma membrane. Error bars are SD. *p* values were calculated from two-tailed Mann-Whitney test.

## Discussion

For this study we developed a novel conditional transmembrane peptide to target the T cell receptor. The design of PITCR involved strategically introducing glutamic acid residues into the transmembrane sequence of the human CD3 ζ chain, as previously described for TYPE7, a rationally designed TM ligand for the EphA2 receptor (Alves et al., 2018). We additionally introduced small and polar residues (**Figure 1-Figure supplementary 1 A**). PITCR selectively inserted into synthetic lipid vesicles at acidic pH (**Figure 1-Figure supplementary 1**). However, we observed that TCR activation in Jurkat cells was severely disrupted by acidic pH (data not shown). We therefore performed experiments at physiological pH, where we observed that PITCR efficiently targeted cells (**Figure 5 and Figure 6**). This observation is not unexpected. The acidity-responsive TYPE7 also targeted cellular membranes at neutral pH because the presence in the membrane of its target receptor shifts the pH-responsiveness to cause membrane insertion at pH 7.4. We suggest that a similar situation might occur for TCR. However, we speculate that PITCR will more effectively inhibit T cells that reside and survive in acidic environments. The solubility of PITCR is a useful property to facilitate delivery to tissues. Furthermore, the pH-responsiveness of PITCR could potentially be used for targeted therapies in pathologies that are characterized by acidic extracellular environments, including inflammatory (Andreev et al., 2007) and autoimmune diseases (Marunaka, 2015), and solid tumors (Cheng et al., 2015). PITCR could also be used to overcome a critical limitation of allogeneic CART therapy, since TCR inhibition is required to prevent graft-versus-host disease resulting from side effects of this therapy. The use of PITCR would additionally overcome the risk resulting from genetic manipulation (i.e., viral transfection) of allogeneic T cells before injection into patients (Michaux et al., 2022).

PITCR caused robust inhibition throughout the signaling cascade that is triggered when TCR is activated, from phosphorylation of the ζ chain to calcium influx. Upon TCR ligation, immunoblot analysis showed that PITCR inhibited phosphorylation at multiple sites in Zap70, LAT, SLP76 and PLCγ1 (**Figure 2**), but not of Lck (**Figure 2-Figure supplementary 2**). The observed specific inhibition of TCR-downstream signaling suggests that PITCR is unlikely to inhibit/activate a broad range of kinases or phosphatases. Rather, PITCR is likely to specifically inhibit TCR triggering, while maintaining Lck association with coreceptors and its phosphorylation (Ashouri et al., 2022; Guy et al., 2022).

What is the molecular mechanism of PITCR inhibition of TCR? A recent molecular dynamics simulation study of TCR (Prakaash et al., 2021) offers clues on a possible binding site for PITCR. The model reveals a hydrophobic gap between the TM domain of the ζ and ε(δ) subunits that can accommodate PITCR (Figure 9). Interestingly, the cryo-EM structure of TCR reveals a binding interface between the TM of ζ and ε(δ) (Dong et al., 2019). PITCR maintains all the residues that form the helical face of ζ that interacts with ε(δ). Therefore, we propose that PITCR binds into the membrane gap between ζ and ε(δ), where it establishes native interactions with the ε(δ) subunit. We found that when we replaced glycine at position 41 with proline, the peptide no longer inhibited calcium influx (**Figure 3**). We speculate that proline induced a local helical kink in PITCR that prevented binding to the TM gap, or a productive interaction between PITCR and the ε(δ) subunit of TCR.

**Figure 9.**
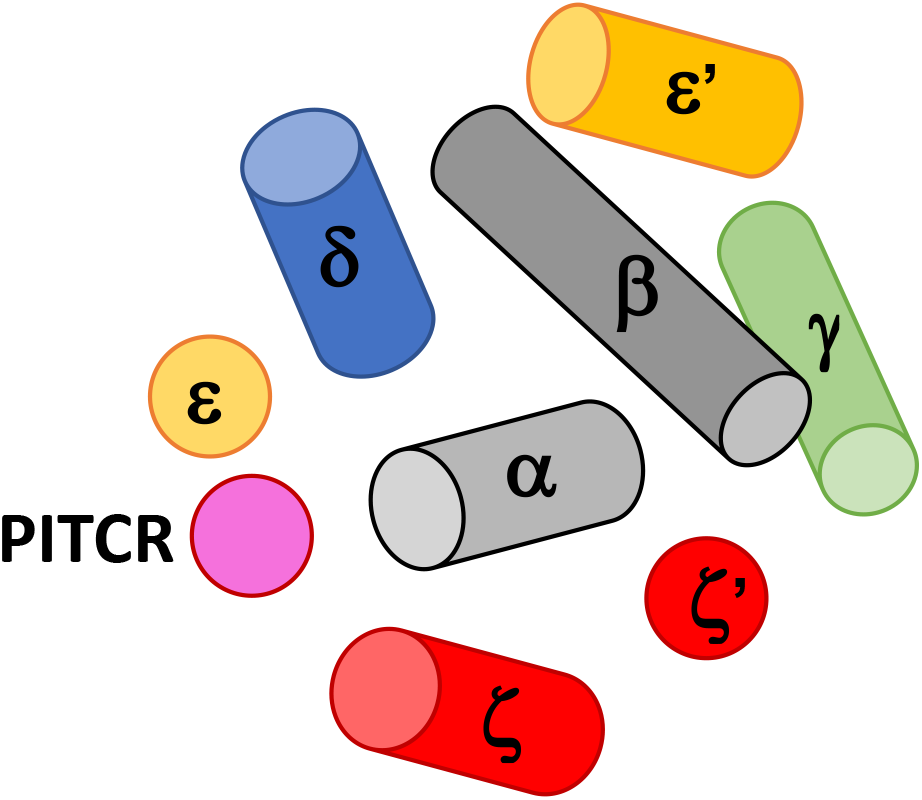
Cartoon model of PITCR interaction with TCR. Schematic view looking from the extracellular side of the TM bundle of TCR, with PITCR is shown in magenta. TM helices are shown as cylinders or circles. The disulfide bond between the ζ subunits is not shown for clarity.

The results of the immunoprecipitation experiments in the presence of DDM (Figure 8) showed that upon TCR activation with OKT3, ζζ strengthened its association with the β subunit. We observed that PITCR caused the opposite effect. This result shows that PITCR acts by reversing the allosteric changes in TM compactness induced by OKT3 activation. However, there was an interesting difference in the interaction with the ε subunit, which increased with OKT3 activation but was not altered by PITCR. This observation suggests that not all the TM interactions are equally important with regards to contributing to TCR activation. Our results additionally suggest that the interface between the ζ and β subunits is a target for pharmacological inhibition of the TCR.

We based the DDM immunoprecipitation assay on a previous report (Lanz et al., 2021), but our protocol contains significant differences. We performed the immunoprecipitation for the native receptor, while the previous protocol involved IP of an HA-tagged TCR, which was also activated by different means to ours. Probably as a result of these differences, our DDM immunoprecipitation result showed an opposite effect into how TCR activation affected the association of TCRαβ with ζζ (Brazin et al., 2018; Lanz et al., 2021). Nevertheless, the two experimental lines of evidence still agree in showing changes in the quaternary stability of the TCR upon activation, and support that this effect could be an important element of the allosteric activation of the TCR. Overall, our data indicate that binding of PITCR to the transmembrane region of TCR results in opposite allosteric changes to those that occur upon TCR activation, and we propose this effect reduces signal transduction into the intracellular region.

Even though the activation of TCR is an intricate process (Chai, 2020; Dong et al., 2019; Mariuzza et al., 2020; Reinherz, 2019), significant progress has been made to understand the conformational changes it entails. For example, in response to TCR engagement, the juxtamembrane domains of the ζζ homodimer have been experimentally reported to switch from a divaricated to a parallel conformation (Lee et al., 2015). TCR activation additionally involves a conformational change of the ITAMs of CD3ε that releases the interaction of basic residues in the intracellular domain of CD3 ζ (Zhang et al., 2011) and ε (Xu et al., 2008) giving access to Lck for phosphorylation. We suggest a plausible mechanism where PITCR insertion into the gap between ζζ homodimer and the rest of TCR-CD3 complex may additionally hinder the activating conformational change in ζζ. This hypothesis might further explain the observed decrease of ζ/β2 in the PITCR-treated Jurkat cells upon TCR activation.

The human adaptive immune system contains numerous types of T cells. T cells present a broad repertoire of TCR (Davis & Bjorkman, 1988). For example, in human peripheral blood samples, 10^4^ varieties of TCRβs can be found (Kidman et al., 2020), while more than 10^15^ combinations of TCRαβ are could theoretically be formed (Davis & Bjorkman, 1988) (Wong et al., 2007) (Freeman et al., 2009). The TCR repertoire plays a critical role in the adaptive immune response, but it also brings challenges to achieve immunosuppression to treat inflammatory and autoimmune diseases. Our data show that PITCR interacts with different types of TCRs: it inhibited TCR signaling in Jurkat-WT (**Figure 1, Figure 2 and Figure 3**), and colocalized with the cSMAC structure formed by TCR(AND) in primary CD4^+^ T cells (**Figure 6**). Furthermore, PITCR reduced CD69 upregulation in OT1-TCR Jurkat cells (**Figure 4**). These results suggest that PITCR may possess the ability to interact with a broad range of TCR types. We hypothesized that this would be possible because the PITCR design is based on targeting the TM region of the TCR, a sequence that is largely conserved.

Taken together, our data show that rational design of a membrane ligand inhibits TCR activation. PITCR has potential clinical value to treat autoimmune and inflammatory diseases or to avoid transplant rejection. The strategy used to design PITCR can be applied to develop targeted ligands for other receptors causative of disease.

## Materials and Methods

### Cell lines

Human male leukemic Jurkat T cells (Clone E6-1, TIB-152) were obtained from the American Type Culture Collection (Manassas, United States). Jurkat.OT1-TCRα-GFP-TCRβ.hCD8^+^ (J.OT1.hCD8^+^) cells and T2-K^b^ cells were kindly provided by Arthur Weiss (UCSF). Cell lines were maintained in RPMI 1640 (Gibco 11875119) supplemented with 10 % fetal bovine serum (Gibco 10437028), 1 % L-Glutamine (Gibco 25030081) and 1 % penicillin & streptomycin (Gibco 15140122) in a 37 °C & 5 % CO_2_ humid tissue culture incubator (Panasonic Healthcare, Wood Dale, United States). The identity of Jurkat cells was authenticated using ATCC services. Mycoplasma contamination was ruled out by PCR (Abcam 289834).

### Peptide synthesis

Peptides were synthesized by Thermo Fisher Scientific (Waltham, United States) and were confirmed by matrix-assisted laser desorption ionization-time-of-flight (MALDI-TOF) mass spectrometry and reverse phase high performance liquid chromatography (HPLC). Purities of the peptides are over 95 %. PITCR sequence: DPKLSYLLDGILFGYGVELTALFLEVGFSESAD.

### Intracellular calcium assay

Jurkat-wild-type (WT) cells were washed twice with PBS and then stained with the calcium sensor dye Indo-1 AM (Invitrogen I1223), at 37 °C and 5 % CO_2_ for 30 mins as described (Lo et al., 2018; Lo et al., 2019). The final concentration of Indo-1 AM was 1 μM. Stained Jurkat cells were washed twice with PBS and then were transferred to a 96 well flat bottom black plate. PITCR was added and incubated at 37 °C and 5 % CO_2_ for 20 mins. Next, the plate was transferred to a prewarmed (37 °C) and 5 % CO_2_ Cytation V plate reader (BioTeK, Winooski, United States) and incubated for another 10 mins. The final concentration of PITCR in the each well was 10 μM. The baseline for each well was recorded for the first 100 seconds, followed by auto-injection of anti-CD3 (OKT3 clone, Tonbo-70-0037). The final concentration of anti-CD3 was 1 μM. Ionomycin (Invitrogen I24222) was used as a positive control to prove that Jurkat cells respond to calcium influx. The fluorescent signal collected from Jurkat-WT cells without staining was subtracted from the signal collected from Indo-1 AM stained cells since Jurkat-WT cells have auto-fluorescent signals.

### SDS-PAGE Immunoblot analysis of proximal signal molecules of TCR-CD3, ζ-Y83 and ζ-Y142

Jurkat-WT cells were washed twice with PBS and treated with PITCR at 37 °C and 5 % CO_2_ for 30 mins, followed by stimulation with anti-CD3-antibody (OKT3 clone) for 5 mins. The final cell density was 5 × 10^6^ cells/ml and the final concentration of PITCR was 10 μM. The final concentration of anti-CD3 was 1 μM. Cells were lysed in 1 % NP-40 lysis buffer (50 mM Tris-Cl pH 7.4, 150 mM sodium chloride, 2 mM PMSF, 5 mM EDTA, 0.25 % sodium deoxycholate with proteinase inhibitors (Thermo Scientific A32955) and phosphatase inhibitors (Sigma-Aldrich P0044) for 30 minutes on ice, followed by centrifugation of 16.2 × 10^3^ g, 30 mins, 4 °C. Supernatants were collected and detergent-compatible protein assay (Bio-Rad5000112) was performed to quantify the protein concentration of each sample. Equal amounts of protein samples were run in 10 %, 12 % or 15 % SDS-PAGE gels and transferred to 0.45 μm nitro cellulose membranes. Membranes were blocked with 3 % bovine serum albumin (BSA) dissolved in TBS, followed by overnight incubation with primary antibodies at 4 °C. IRDye 800CW or IRDye 680LT secondary antibodies were used to incubate the blots at the 2^nd^ day, followed by detection with an Odyssey Infrared Scanner (Li-Cor Biosciences, Lincoln, United States). All primary antibodies were diluted in 5 % BSA dissolved in 0.1 % TBST except specific mentions and all secondary antibodies were diluted in 5 % non-fat milk dissolved in 0.1 % TBST unless mentioned otherwise.

### Co-localization assay and analysis

All steps were performed at room temperature (RT), unless noted otherwise. Jurkat-WT cells were washed once with PBS, resuspended in RMPI1640 phenol-red free media, and treated with dylight680 labeled PITCR (PITCR680) at 37 °C and 5 % CO_2_ for 15 mins, followed by stimulation in presence or absence of anti-CD3 (OKT3 clone) for 5 mins. The final concentration of anti-CD3 was 1 μM. The final cell density was 5 × 10^6^ cells/ml and the final concentration of PITCR680 was 10 μM. PITCR680 treated cells were washed twice with cold PBS and resuspended in cold RPMI1640 phenol-red free media. 100 μL of cells were transferred to each chamber of μ-Slide 8 Well ibiTreat (ibidi80826) and rested for 10 mins. Next, RPMI1640 phenol-red free media was removed gently. 150 μL cold fixing buffer (1.998 % - formaldehyde and 0.2 % - glutaraldehyde in filtered PBS) was added and incubated for 10 mins. Fixed PITCR680 treated Jurkat cells were washed twice with cold DPBS/Modified (HycloneSH30264.01). 150 μL permeabilization buffer (0.1 % Triton X100 in PBS) was incubated with the sample for another 15 mins. Each chamber was washed with blocking buffer (5 % goat serum in 0.01 % PBST) twice. The sample was blocked 1 hr, followed by washing once with antibody dilution buffer (1 % BSA in 0.01 % PBST). The samples were incubated with 1:200 diluted anti-CD3ε (UCHT1 clone, sc-1179) primary antibody in the wet box at4 °C, overnight. On the 2^nd^ day, each chamber was washed twice with cold DPBS/Modified (HycloneSH30264.01). 1:200 diluted secondary antibody (Invitrogen A32723) was incubated with samples in a foil covered wet box for 1 hr, followed by washing twice with cold DPBS/Modified (HycloneSH30264.01). 1:1000 diluted DAPI (Thermo Scientific 62248) was stained with samples for 2 mins, followed by washing once with cold DPBS/Modified. The samples were mounted with Vectashield (H-1000), followed by sealing the chambers with parafilm until imaging.

Samples were imaged using a Leica SP8 White Light Laser Confocal Microscope (Leica, Wetzlar, Germany) equipped with a 63X oil immersion objective, zoom 5 through LAS X software. Z-stack scanning was applied, followed by a lightning deconvolution analysis. Each image was chosen from one time point at each Z-stack section. Graphic profile curves of the Region of Interest in Zoom-in images were analyzed using Image J RGB Profile Plot Plugin. The Mander’s M1 and Pearson’s *r* coefficients were calculated using Image J Just Another Colocalization Plugin (JACoP).

### CD69 activation flow cytometry assay

2×10^6^ cells/ml T2-K^b^ cells were washed twice with PBS and treated with a series of diluted ovalbumin (OVA) derived peptide (SIINFEKL) at 37 °C and 5 % CO_2_ for 1 hr. For this assay we used J.OT1.hCD8^+^ cells, which are engineered human Jurkat T cells, as described (Lo et al., 2018). 5 × 10^6^ cells/ml J.OT1.hCD8^+^ cells were washed twice with PBS and treated with PITCR at 37 °C and 5 % CO_2_ for 30 mins. PITCR-treated J.OT1.hCD8^+^ cells were added to OVA-treated T2-K^b^ cells and the ratio of these two cells was 1:1. Final concentration of PITCR was 10 μM. The incubation time was 3 hrs. Next, anti-CD69 - Allophycocyanin (Biolegend 310910) and Isotype - Allophycocyanin (Biolegend 400122) were applied to stain the cells, respectively followed by LSRII Flow Cytometer (BD Bioscience, Franklin Lakes, United States) analysis. Data was quantified using FlowJo_v10.8.0 software.

### Co-Immunoprecipitation of TCR-CD3 complex and immunoblot analysis

Jurkat-WT T cells were washed twice with PBS and treated with Dylight680 labeled PITCR at 37 °C and 5 % CO_2_ for 30 mins, followed by stimulation with 1 μM anti-CD3 antibody (OKT3 clone). Cells were lysed in 2.5 % Diisobutylene Maleic Acid (DIBMA, AnatraceBMA101) lysis buffer (20 mM Tris-Cl pH 8.0, 137 mM NaCl, 2 mM EDTA, 1 mM PMSF, 5 mM iodoacetamide, 1 mM NaF, proteinase inhibitors (Promega G6521) and phosphatase inhibitors (Sigma-Aldrich P0044 and P5726) at warm room (37 °C) for 2 hrs followed by rotating at least 12 hrs in the cold room (4 °C). Lysates were ultracentrifuged 10,000 g, 4 °C, 45 mins to get rid of debris, and the supernatants were collected and ultracentrifuged again 10,000 g, 4 °C, 1hr. The protein concentrations were quantified using a detergent compatible protein assay. 40 μl whole lysates were saved to detect the TCR-CD3 complex. 1 % BSA blocked protein G agarose (Thermo Scientific 20398) and anti-CD3ε (UCHT1 clone, sc-1179) were added to the rest of lysates. Samples were gently rotated at cold room (4 °C) for 16 hrs. Samples were centrifuged 5,000 g, 4 °C, 3mins, followed by washing twice with cold wash buffer (20 mM Tris-Cl pH 8.0, 137 mM NaCl, 2 mM EDTA, 1 mM PMSF, 5 mM iodoacetamide, 1 mM NaF) and rinsing once with cold wash buffer. SDS sample buffer was applied to elute the captured protein followed by incubation at 95 °C for 5 mins.

Equal amount of whole lysate samples and captured protein samples were loaded in 15 % SDS-PAGE glycine gels and transferred to 0.45 μm or 0.2 μm nitro cellulose membranes. 3 % BSA was used to block the membranes, followed by incubation with primary antibodies: anti-TCRβ, anti-CD3ε and anti-CD3ζ (6B10.2 clone) overnight at 4 °C. In the whole lysates’ group, the housekeeping protein β-actin was also probed. PITCR was directly detected using channel 700 nm of an Odyssey Infrared Scanner (Li-Cor Biosciences, Lincoln, United States). The 2^nd^ day, same approaches were performed as described in SDS-PAGE Immunoblot analysis of proximal signal molecules of TCR-CD3, ζ-Y83 and ζ-Y142.

### Liposome preparation

1-palmitoyl-2-oleolyl-glycero-3-phosphocholine (POPC) and 1-palmitoyl-2-oleoyl-sn-glycero-3-phospho-L-serine (POPS) were purchased from Avanti Polar Lipids, Alabaster, AL. POPC and POPS were dissolved in cold chloroform and stocks prepared at 32.89 mM. POPS and POPC were mixed in a round-bottom test tube, dried together under a stream of argon gas, and placed in a vacuum overnight. The dried lipid film was rehydrated in 10 mM sodium phosphate buffer (pH 7.4), followed by extrusion with a Mini-Extruder (Avanti Polar Lipids, Alabaster, AL) through a 0.1 μm Nuclepore Track-Etch membrane (Whatman, Maidstone, United Kingdom). The final large unilamellar vesicles (LUVs) stock concentration was 4 mM. LUVs contain 10% POPS and 90% POPC.

### Circular dichroism (CD)

CD experiments were performed as described previously (Nguyen et al., 2015). Briefly, the secondary structure of PITCR in aqueous solution (10 mM sodium phosphate, pH 7.4) and liposomes (10 % POPS and 90 % POPC LUVs at pH 5.0 and pH 7.4, respectively) was determined in a Jasco J-815 spectropolarimeter at RT. The Circular Dichroism (CD) spectra were measured from 195 nm to 260 nm in a 2 mm path length quartz cuvette. The peptide to lipid molar ratio was 1: 200 and the final concentration of the peptide was 5 μM. To obtain the desired pH, the experimental samples were adjusted by adding either 100 mM sodium phosphate (pH 7.4) or 100 mM sodium acetate (pH 4.0). The appropriate liposome or buffer backgrounds were subtracted. Molar ellipticity was calculated with the following equation: [*θ*] = *θ*/[10*lc*(*N* – 1)], where *θ* is the measured ellipticity in millidegree, *l* is the cell path length, *c* is the protein concentration and *N* is the number of amino acids (*N*= 33).

### pK_CD_ determination assay

The apparent pK _CD_ is defined as a pH midpoint, where half of the peptide changes its secondary structure in presence of liposomes (Scott et al., 2017). The liposome preparation (10 % POPS and 90% POPC LUVs) was followed as described in liposome preparation. To reach a series of different desired pH values, the experimental samples were adjusted with either 100 mM sodium phosphate or 100 mM sodium acetate. The final pH was measured by a 2.5-mm bulb pH electrode (Microelectrodes, Bedford, NH) after recording CD spectrum. The CD spectra were recorded from 220 nm to 262 nm. The appropriate liposome blanks were subtracted. The ellipticity values at 222 nm were subtracted from that at 262 nm. The subtracted ellipticity were plotted for a range of pH values. The pK _CD_ was fitted in the following equation: 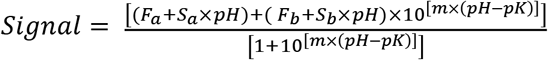, where *F_a_* is the acidic baseline, *F_b_* is the basic baseline, m is the slope of the transition, and pK is the midpoint of the curve.

### Peptide conjugation

Cysteine was added to the N-terminal of PITCR, termed NEC-PITCR (sequence: ECDPKLSYLLDGILFGYGVELTALFLEVGFSESAD). NEC-PITCR was labeled with dylight680 maleimide (Thermo Scientific-46618) and AZDye 555 maleimide (Fluoroprobes-1168-1). Both dyes labeled peptides were purified using reverse phase HPLC purification to remove unconjugated dye. The molecular weight was confirmed by MALDI-TOF. After that, samples were aliquoted, lyophilized and stored at −80 °C.

### MALDI-TOF mass spectrometry

PITCR associated peptides were dissolved in 1 mM sodium phosphate buffer (pH 7.4, filtered). The matrix α-Cyano-4-hydroxycinamic acid (TCI C1768), was dissolved in 75 % HPLC-level acetonitrile coupled with 0.1 % TFA and sonicated 15 minutes, RT. The dissolved samples were mixed with the dissolved matrix. After that, the mixed matrix samples were loaded onto the MSP target plate (Bruker, Billerica, United States) drop by drop and dried using filtered air. The Bruker Microflex MALDI-TOF mass spectrometer (Bruker, Billerica, United States) was calibrated with ProteoMass MALDI-MS calibration standards (Sigma-Aldrich I6279-5X1VL, I6154-5X1VL, C8857-5X1VL and P2613-5X1VL). All PITCR associated peptides were measured in a negative mode. The pHLIP was measured in a positive mode. Data were analyzed using FlexAnalysis software (Bruker, Billerica, United States) and graphs were plotted using Origin 9.1 (research lab) software.

### HPLC

All peptides were dissolved in 1 mM sodium phosphate buffer (pH 7.4, filtered). Dissolved peptides were injected into an Agilent 1200 series HPLC system (Agilent Technologies, Santa Clara, United States). A semi-preparative Agilent Zorbax 300 SB-C18 column (P.N. 880995-202) was used to purify the PITCR associated peptides. A stable-bond analytical Agilent Zorbax 300 SB-C18 column (P.N. 880995-902) was used to identify the purity of each peptide. The running procedure used a gradient (solvent A: 0.05% TFA HPLC-level water plus solvent B: 0.05% TFA HPLC-level acetonitrile) from 5% B to 100% B. PITCR associated peptides were eluted around 80% B.

### Immunoprecipitation and immunoblot analysis

Jurkat-WT T cells were washed twice with PBS and treated with PITCR at 37 °C and 5 % CO_2_ for 30 mins, followed by stimulation with 1 μM anti-CD3 antibody (OKT3 clone). Cells were lysed in cold 0.5% Dodecyl-β-D-maltopyranoside (DDM, VWR-97063-172) lysis buffer (20 mM Tris-Cl pH 8.0, 10 mM NaF, 166.67 mM NaCl, 20 mM iodoacetamide, Benzonase endonuclease 50 U/ml (Sigma-1016970001), proteinase inhibitors (Thermo Scientific A32955) and phosphatase inhibitors (Sigma-Aldrich P0044 and P5726)) for 30 minutes on ice, followed by centrifugation of 16.2 × 10^3^ g, 15 mins, 4 °C. Supernatants were collected and detergent compatible protein assay was performed to quantify the protein concentration of each sample. 30-50 μl whole lysates were saved to detect the TCR complex. Protein G agarose and anti-CD3ε (UCHT1 clone) were used as described in the methodology of co-IP of TCR. Samples were continuously rotated at 4 °C for 4 hrs. Other steps follow the methodology of co-IP of TCR, except the wash buffer (20 mM Tris-Cl pH 8.0, 10 mM NaF, 166.67 mM NaCl, 20 mM iodoacetamide, Benzonase endonuclease 50 U/ml, Sigma-1016970001) and the elution condition (70 °C for 10 mins). Immunoblot analysis were performed as described in the methodology of co-IP of TCR.

### Antibodies

| Antibodies | Sources | Catalogue # | Dilutions |
| --- | --- | --- | --- |
| Zap70 (pY319) | Cell Signaling Technology | 2717 | 1:1000 (WB) |
| Zap70 (pY493) | Cell Signaling Technology | 2704 | 1:1000 (WB) |
| Zap70 (total) | Cell Signaling Technology | 3165 | 1:1000 (WB) |
| LAT (pY191) | Cell Signaling Technology | 3584<br>(discontinued) | 1:1000 (WB) |
| LAT (pY132) | Abcam | ab4476 | 1:2000 (WB) |
| LAT total | Cell Signaling Technology | 45533 | 1:1000 (WB) |
| Lck (pY394) | R & D systems | 755103 | 1:2000 (WB) |
| Lck (pY505) | Cell Signaling Technology | 2751 | 1:1000 (WB) |
| Lck (total) | Cell Signaling Technology | 2984 | 1:1000 (WB) |
| β-actin | Abcam | ab6276 | 1:5000 (WB) |
| CD3 ε (OKT3) | Tonbo biosciences | 70-0037 | 1:50 (stimulation) |
| CD3 ε (UCHT1) | Santa Cruz biotechnology | sc-1179 | 1: 200 (IF), IP<br>(see methodology) |
| CD3 ε (CD3-12) | Cell Signaling Technology | 4443 | 1:1000 (WB) |
| ζ (6B10.2) | Santa Cruz biotechnology | sc-1239 | 1:500 (WB) |
| $\zeta$ (pY142) | BD Biosciences | 558402 | 1:1000 (WB) |
| $\zeta$ (pY83) | Abcam | ab68236 | 1:1000 (WB) |
| TCR $\beta$ | Cell Signaling Technology | 77046 | 1:2000 (WB) |
| SLP76 (pY128) | BD Biosciences | 558367 | 1:2000 (WB) |
| PLCγ1 (pY783) | Cell Signaling Technology | 2821 | 1:1000 (WB) |
| IgG | Cell Signaling Technology | 5415 | IP (see methodology) |
| CD69-APC (FN50) | Biolegend | 50-166-584 | 1:100<br>(Flowcytometry) |
| IgG-APC | Biolegend | 50-168-838 | Flowcytometry |
| Goat anti-Mouse IgG (H+L)<br>Highly Cross-Adsorbed<br>Secondary Antibody, Alexa<br>Fluor™ Plus 488 | ThermoFisher Scientific<br>(Invitrogen) | A32723 | 1: 200 (IF) |
| IRDye® 800CW Goat anti-Rabbit IgG Secondary Antibody | LI-COR Bioscience | 926-32211 | 1:5000 (WB) |
| IRDye® 800CW Goat anti-Mouse IgG Secondary Antibody | LI-COR Bioscience | 926-32210 | 1:5000 (WB) |
| IRDye® 800CW Goat anti-Rat<br>IgG Secondary Antibody | LI-COR Bioscience | 925-32219 | 1:5000 (WB) |

### TIRF imaging of AND-TCR T cells

AND-TCR T cells were incubated with PITCR-AZDye555 by mixing 50 μL of a 100 μM solution in 10 mM sodium phosphate buffer pH 7.4 with 450 μL of 2.5 M cells/mL T cells in RVC medium with IL-2 (final concentration of 10 μM PITCR-AZDye555, 2.2 M cells/mL, 37 °C, 30 min), rinsed by imaging buffer, then applied to the imaging chamber with SLB functionalized with ICAM-1 (~20 molecules/μm2) and pMHC (19-23 molecules/μm2, MCC peptide 9.1% labeled with Atto647N) at 37 °C. The real-time images of just-adhering cells with RICM, TIRF at 561 nm excitation (50 ms exposure), and TIRF at 640 nm excitation (500 ms exposure) channels were recorded at the average frame rate of 1 frame per 4 seconds. After 15-30 min, the snapshots of the cells forming immune synapses were recorded with the same three channels. The experiment was performed with 5 cells (real-time) and about 50-100 cells (snapshots) from 1 mouse. The cells with dominant plasma-membrane-bound PITCR signal could be found only in snapshot measurements due to low population. The vehicle control was performed by treating the cells with phosphate buffer instead of PITCR-AZDye555 solution using the cells from the same mouse.

#### Reagents

T cell culture medium: DMEM (Gibco, Thermo Fisher) with 10% FBS, 1 mM sodium pyruvate, 2 mM L-glutamine, 1× Corning nonessential amino acids (Fisher Scientific), 1x Corning MEM vitamin solution (Fisher Scientific), 0.67 mM L-arginine, 0.27 mM L-asparagine, 14 μM folic acid, 1x Corning Penicillin/Streptomycin (100 IU, 0.1 mg/mL respectively) (Fisher Scientific), 50 μM β-mercaptoethanol.

Imaging buffer for TIRF experiment: 20 mM HEPES, 137 mM NaCl, 5 mM KCl, 1 mM MgCl_2_, 1.8 mM CaCl_2_, 0.1% w/v D-glucose, 0.1% w/v BSA, pH 7.4.

#### AND-TCR T cell culture

Primary AND-TCR T cells were prepared and cultured basically as previously described (Smith et al., 2011). T cells from the lymph nodes and spleens were harvested from (B10.Cg-Tg(TcrAND)53Hed/J) × (B10.BR-H2k2 H2-T18a/SgSnJ) hybrid mice (Jackson Laboratory) and kept in T cell culture medium (day 1). The cells were activated by 2 μM MCC peptide at day 1, then cultured with IL-2 after day 2. Cells were used at day 5 or 6 for imaging. All animal work was performed with prior approval by Lawrence Berkeley National Laboratory Animal Welfare and Research Committee under the approved protocol 177003.

#### pMHC and ICAM-1 preparation

ICAM-1 extracellular domain with His10 tag and MHC class II I-Ek with two His6 tags were expressed and purified as previously described (Nye & Groves, 2008).

Peptides for pMHC were prepared and loaded to MHC molecule as previously described (O’Donoghue et al., 2013) MCC peptide (ANERADLIAYLKQATK) and MCC-GGSC (ANERADLIAYLKQATKGGSC) were synthesized on campus (D. King, Howard Hughes Medical Institute Mass Spectrometry Laboratory at University of California, Berkeley) or commercially (Elim Biopharmaceuticals, Hayward, CA). MCC-GGSC was labeled with Atto647N-maleimide (ATTO-TEC), purified by C18 column reversed phase HPLC, and identified by MALDI-TOF mass spectrometry.

Excess amount of MCC and MCC-GGSC-Atto647N were separately loaded on MHC molecules in loading buffer (PBS acidified with citric acid to pH 4.5, 1% BSA) over night at 37°C. Then mixed at a 10:1 molar ratio to achieve 9.1% labeling efficiency. The mixture was diluted-concentrated with TBS and 10 kDa MWCO filters (Spin-X UF, Corning) for two times to remove excess peptides, then used for bilayer functionalization.

#### Supported lipid bilayer preparation

Small unilamellar vesicles with 98 mol% 1,2-dioleoyl-sn-glycero-3-phosphocholine (DOPC, Avanti Polar Lipids) and 2 mol% 1,2-dioleoyl-sn-glycero-3-[(N-(5-amino 1-carboxypentyl)iminodiacetic acid)succinyl] nickel salt (DGS-NTA, Avanti Polar Lipids) were prepared by sonicating a 0.5 mg/mL lipid suspension in water followed by centrifugation (21,000 *g*, 20 min, 4°C). Then, supported lipid bilayer (SLB) was prepared upon 25 mm #1.5 glass coverslip set into Attofluor cell chamber (Invitrogen, Thermo Fisher). Coverslips were cleaned by sonication in 1:1 water:2-propanol then etched with piranha solution (1:3 mixture of 30% H_2_O_2_ and sulfuric acid), rinsed by water and set into clean chambers. SLB was formed by adding 1:1 mixture of SUV solution and TBS into chambers and incubating for more than 30 min. SLB were rinsed with TBS then incubated with 10 mM NiCl_2_ in TBS for 5 min. Chambers were then incubated in imaging buffer for more than 30 min for blocking defects by BSA, then used for functionalization. ICAM-1 (~10 nM) and pMHC (concentration adjusted by determined densities) were added to chambers and incubated for 30 min, then rinsed by imaging buffer. The ICAM-1 density is estimated to be ~20 molecule/μm^2^ based on a previously reported estimate (Lin et al., 2019). pMHC labeled by Atto647N was imaged by TIRF to determine the molecular density. The density was determined by extrapolating the calibration curve of density-intensity relationship from lower densities where the molecular density can be directly determined by single molecule localization (below 0.5 molecules/μm^2^) using TrackMate (Tinevez et al., 2017).

#### TIRF microscopy and image processing

TIRF microscopy was performed on a motorized inverted microscope (Nikon Eclipse Ti-E; Technical Instruments, Burlingame, CA) with Lumen Dynamics X-Cite® 120LED Fluorescence Illumination System (Excelitas Technologies, Waltham, MA, USA) and a motorized stage (MS-2000; Applied Scientific Instrumentation, Eugene, OR). A laser launch with 561-, and 640-nm diode lasers (Coherent OBIS, Santa Clara, CA) was aligned into a custom-built fiber launch (Solamere Technology Group Inc., Salt Lake City, UT). For TIRF imaging, laser excitation was illuminated through a four bands beam splitter (ZT405/488/561/640rpc) to the objective lens (NA 1.49, 100×, oil immersion, Apochromat TIRF, Nikon), then filtered through an emission filter (ET600/50M or ET700/75M). For RICM, LED excitation was illuminated through D546/10x excitation filter and 50/50 beam splitter. Emission was captured on an EM-CCD (iXon Ultra 897; Andor Inc., South Windsor, CT). All optical filters were purchased from Chroma Technology Corp. (Bellows Falls, VT). The sample and objective lens were kept at 37 °C with temperature controller system (CU-109, Live Cell Instrument, Republic of Korea). The equipment was controlled using the software MicroManager (Edelstein et al., 2010) Laser power and exposure time was set to 2 mW, 50 ms for 561 nm excitation and 1 mW, 500 ms for 640 nm excitation. The pixel size was 0.16 μm square and the field of view was 81.92 μm square (512×512 pixels).

Cell footprint was determined from RICM images with the following procedures. RICM image was gaussian-blurred (sigma: 2 pixels), manually background-subtracted, and converted to absolute values pixel-wise. The obtained intensity images were segmented by semi-automatic way: the inner region and the encompassing region of the cell of interest were manually selected. Image was thresholded by the intensity in the inner region multiplied by an arbitrary factor of 0.5. The obtained segment was filtered within the encompassing region, then cleaned by binary opening (kernel: 3×3 pixels square). The regions smaller than 200 pixels were deleted and the remaining regions (multiple regions were allowed to exist if any) were used as the cell footprint.

The background signal was measured using the chamber containing only imaging buffer and subtracted from TIRF images. The inhomogeneity of the TIRF illumination were corrected using the images from the solution of rhodamine B (561 nm excitation, from Sigma Aldrich) and 3,3’-diethylthiadicarbocyanine iodide (640 nm excitation, from Sigma Aldrich).

### NFAT activation assay

To assay activation of AND-TCR primary murine CD4+ T cells, cells were transduced with a LAT-eGFP-P2A-NFAT-mCherry bicistronic construct on day 3 of primary cell culture as previously described (Smith et al., 2011). Cells were assayed on day 5. All animal work was performed with prior approval by Lawrence Berkeley National Laboratory Animal Welfare and Research Committee under the approved protocol 177003.

For each imaging chamber, 2.5 million cells were resuspended to 5 million/mL in 450 μL imaging buffer and 50 μL of 100 μM unlabeled PITCR in 10 mM sodium phosphate buffer pH 7.4. Control samples were treated the same, with PITCR omitted. Cells were incubated for 30 min at 37 °C and then directly added to SLBs functionalized with ICAM and pMHC in Attofluor chambers containing 500 μL imaging buffer equilibrated to 37 °C. Cells interacted with the bilayer for 20 min before acquiring snapshots to analyze for NFAT activation state. Cells transduced with reporter proteins were identified using the LAT signal in the 488 TIRF channel to minimize bias in which cells were imaged and subsequently analyzed. Single sets of RICM, 488 TIRF, and 561 epifluorescence images were taken for at least 30 fields of view and at least 50 live cells 20-50 min after adding cells to the SLB. Three z-positions were acquired for 561 epifluorescence at 0, 3, and 6 μm above the TIRF plane in order to clearly resolve each cell’s nucleus and the distribution of NFAT-mCherry between the cytoplasm and nucleus. Only cells with substantial contact with the bilayer, as defined by the RICM footprint, were included in the analysis of the fraction of activated cells. Cells were defined as active if the NFAT-mCherry signal in the nucleus was equal to or greater than the signal in the cytoplasm, assessed manually, indicating that the NFAT-mCherry reporter protein was translocating to the nucleus. The fraction of activated cells was determined for each bilayer density and pre-treatment condition and error bars denote the standard error of the mean. The density of pMHC on each bilayer was determined before the addition of cells. Snapshots of pMHC in at least 20 fields of view were taken in the 640 TIRF channel at 20 mW power at the source and 20 ms exposure time. Particles were counted using TrackMate and density was determined as the particle count divided by the total area of all fields of view. A calibration curve relating density and intensity was used to measure the density of high-density bilayers for which single particles are not able to be resolved (~ 0.7 μm^-2^)

### Statistical analysis

All statistical analyses of experiments were performed using GraphPad Prism 9.4.0. P values are provided as exact values. 95% confidence level was used to determine statistically significance in all experiments and ns stands for not significant. All statistical correspond to biological replicates only and all n values reflect biological replicates. Detailed statistical analyses were illustrated in each result.

## Supporting information

Full supplementary materials

## Acknowledgements

This work was supported by grant R35GM140846 (to F.N.B.), and by a Faculty-Graduate student award to YYJ (University of Tennessee). We thank Art Weiss (UCSF) for generous advice to YYJ and for providing J.OT1.CD8 and T2Kb cells, and to L. Teyton (Scripps Research) and M. Davis (Stanford University) for providing the MHC and ICAM-1 bacmids. We also appreciate the advice provided to YYJ by Barry Bruce (University of Tennessee), Matthew Call (Walter and Eliza Hall Institute of Medical Research), and Peiqing Sun (Wake Forest Baptist Medical Center). We are also thankful for the technical advice of Tim Sparer and Trevor Hancock (University of Tennessee), Jaydeep Kolape (AMIC, University of Tennessee) and Ed Wright (BRF, University of Tennessee). We appreciate the members of the Barrera lab Jen Schuster, Jordan Pyron and Boomer Russell for insights on the manuscript.

## References

Alves, D. S., Westerfield, J. M., Shi, X., Nguyen, V. P., Stefanski, K. M., Booth, K. R., Kim, S., Morrell-Falvey, J., Wang, B. C., Abel, S. M., Smith, A. W., & Barrera, F. N. (2018). A novel pH-dependent membrane peptide that binds to EphA2 and inhibits cell migration. Elife, 7. https://doi.org/10.7554/eLife.36645

Andreev, O. A., Dupuy, A. D., Segala, M., Sandugu, S., Serra, D. A., Chichester, C. O., Engelman, D. M., & Reshetnyak, Y. K. (2007). Mechanism and uses of a membrane peptide that targets tumors and other acidic tissues in vivo. Proc Natl Acad Sci U S A, 104(19), 7893–7898. https://doi.org/10.1073/pnas.0702439104

Ashouri, J. F., Lo, W. L., Nguyen, T. T. T., Shen, L., & Weiss, A. (2022). TZAP70, too little, too much can lead to autoimmunity. Immunol Rev, 307(1), 145–160. https://doi.org/10.1111/imr.13058

Au-Yeung, B. B., Shah, N. H., Shen, L., & Weiss, A. (2018). ZAP-70 in Signaling, Biology, and Disease. Annu Rev Immunol, 36, 127–156. https://doi.org/10.1146/annurev-immunol-042617-053335

Biswas, K. H., & Groves, J. T. (2019). Hybrid Live Cell-Supported Membrane Interfaces for Signaling Studies. Annu Rev Biophys, 48, 537–562. https://doi.org/10.1146/annurev-biophys-070317-033330

Brazin, K. N., Mallis, R. J., Boeszoermenyi, A., Feng, Y., Yoshizawa, A., Reche, P. A., Kaur, P., Bi, K., Hussey, R. E., Duke-Cohan, J. S., Song, L., Wagner, G., Arthanari, H., Lang, M. J., & Reinherz, E. L. (2018). The T Cell Antigen Receptor alpha Transmembrane Domain Coordinates Triggering through Regulation of Bilayer Immersion and CD3 Subunit Associations. Immunity, 49(5), 829–841 e826. https://doi.org/10.1016/j.immuni.2018.09.007

Bromley, S. K., Burack, W. R., Johnson, K. G., Somersalo, K., Sims, T. N., Sumen, C., Davis, M. M., Shaw, A. S., Allen, P. M., & Dustin, M. L. (2001). The immunological synapse. Annu Rev Immunol, 19, 375–396. https://doi.org/10.1146/annurev.immunol.19.1.375

Call, M. E., Pyrdol, J., Wiedmann, M., & Wucherpfennig, K. W. (2002). The organizing principle in the formation of the T cell receptor-CD3 complex. Cell, 111(7), 967–979. https://doi.org/10.1016/s0092-8674(02)01194-7

Call, M. E., Schnell, J. R., Xu, C., Lutz, R. A., Chou, J. J., & Wucherpfennig, K. W. (2006). The structure of the zetazeta transmembrane dimer reveals features essential for its assembly with the T cell receptor. Cell, 127(2), 355–368. https://doi.org/10.1016/j.cell.2006.08.044

Chai, J. (2020). Atomic-resolution view of complete TCR-CD3 revealed. Protein Cell, 11(3), 158–160. https://doi.org/10.1007/s13238-019-00677-7

Chakraborty, A. K., & Weiss, A. (2014). Insights into the initiation of TCR signaling. Nat Immunol, 15(9), 798–807. https://doi.org/10.1038/ni.2940

Chen, Y., Zhu, Y., Li, X., Gao, W., Zhen, Z., Dong, Huang, B., Ma, Z., Zhang, A., Song, X., Ma, Y., Guo, C., Zhang, F., & Huang, Z. (2022). Cholesterol inhibits TCR signaling by directly restricting TCR-CD3 core tunnel motility. Mol Cell, 82(7), 1278–1287 e1275. https://doi.org/10.1016/j.molcel.2022.02.017

Cheng, C. J., Bahal, R., Babar, I. A., Pincus, Z., Barrera, F., Liu, C., Svoronos, A., Braddock, D. T., Glazer, P. M., Engelman, D. M., Saltzman, W. M., & Slack, F. J. (2015). MicroRNA silencing for cancer therapy targeted to the tumour microenvironment. Nature, 518(7537), 107–110. https://doi.org/10.1038/nature13905

Costes, S. V., Daelemans, D., Cho, E. H., Dobbin, Z., Pavlakis, G., & Lockett, S. (2004). Automatic and quantitative measurement of protein-protein colocalization in live cells. Biophys J, 86(6), 3993–4003. https://doi.org/10.1529/biophysj.103.038422

Courtney, A. H., Amacher, J. F., Kadlecek, T. A., Mollenauer, M. N., Au-Yeung, B. B., Kuriyan, J., & Weiss, A. (2017). A Phosphosite within the SH2 Domain of Lck Regulates Its Activation by CD45. Mol Cell, 67(3), 498–511 e496. https://doi.org/10.1016/j.molcel.2017.06.024

Courtney, A. H., Lo, W. L., & Weiss, A. (2018). TCR Signaling: Mechanisms of Initiation and Propagation. Trends Biochem Sci, 43(2), 108–123. https://doi.org/10.1016/j.tibs.2017.11.008

Davis, M. M., & Bjorkman, P. J. (1988). T-cell antigen receptor genes and T-cell recognition. Nature, 334(6181), 395–402. https://doi.org/10.1038/334395a0

Dong Zheng, L., Lin, J., Zhang, B., Zhu, Y., Li, N., Xie, S., Wang, Y., Gao, N., & Huang, Z. (2019). Structural basis of assembly of the human T cell receptor-CD3 complex. Nature, 573(7775), 546–552. https://doi.org/10.1038/s41586-019-1537-0

Edelstein, A., Amodaj, N., Hoover, K., Vale, R., & Stuurman, N. (2010). Computer control of microscopes using microManager. Curr Protoc Mol Biol, Chapter 14, Unit14–20. https://doi.org/10.1002/0471142727.mb1420s92

Freeman, J. D., Warren, R. L., Webb, J. R., Nelson, B. H., & Holt, R. A. (2009). Profiling the T-cell receptor beta-chain repertoire by massively parallel sequencing. Genome Res, 19(10), 1817–1824. https://doi.org/10.1101/gr.092924.109

Ganti, R. S., Lo, W. L., McAffee, D. B., Groves, J. T., Weiss, A., & Chakraborty, A. K. (2020). How the T cell signaling network processes information to discriminate between self and agonist ligands. Proc Natl Acad Sci U S A, 117(42), 26020–26030. https://doi.org/10.1073/pnas.2008303117

Grakoui, A., Bromley, S. K., Sumen, C., Davis, M. M., Shaw, A. S., Allen, P. M., & Dustin, M. L. (1999). The immunological synapse: a molecular machine controlling T cell activation. Science, 285(5425), 221–227. https://doi.org/10.1126/science.285.5425.221

Guy, C., Mitrea, D. M., Chou, P. C., Temirov, J., Vignali, K. M., Liu, X., Zhang, H., Kriwacki, R., Bruchez, M. P., Watkins, S. C., Workman, C. J., & Vignali, D. A. A. (2022). LAG3 associates with TCR-CD3 complexes and suppresses signaling by driving co-receptor-Lck dissociation. Nat Immunol, 23(5), 757–767. https://doi.org/10.1038/s41590-022-01176-4

He, L., Steinocher, H., Shelar, A., Cohen, E. B., Heim, E. N., Kragelund, B. B., Grigoryan, G., & DiMaio, D. (2017). Single methyl groups can act as toggle switches to specify transmembrane Protein-protein interactions. Elife, 6. https://doi.org/10.7554/eLife.27701

Kidman, J., Principe, N., Watson, M., Lassmann, T., Holt, R. A., Nowak, A. K., Lesterhuis, W. J., Lake, R. A., & Chee, J. (2020). Characteristics of TCR Repertoire Associated With Successful Immune Checkpoint Therapy Responses. Front Immunol, 11, 587014. https://doi.org/10.3389/fimmu.2020.587014

Kuhns, M. S., & Davis, M. M. (2008). The safety on the TCR trigger. Cell, 135(4), 594–596. https://doi.org/10.1016/j.cell.2008.10.033

Lanz, A. L., Masi, G., Porciello, N., Cohnen, A., Cipria, D., Prakaash, D., Balint, S., Raggiaschi, R., Galgano, D., Cole, D. K., Lepore, M., Dushek, O., Dustin, M. L., Sansom, M. S. P., Kalli, A. C., & Acuto, O. (2021). Allosteric activation of T cell antigen receptor signaling by quaternary structure relaxation. Cell Rep, 36(2), 109375. https://doi.org/10.1016/j.celrep.2021.109375

Lee, M. S., Glassman, C. R., Deshpande, N. R., Badgandi, H. B., Parrish, H. L., Uttamapinant, C., Stawski, P. S., Ting, A. Y., & Kuhns, M. S. (2015). A Mechanical Switch Couples T Cell Receptor Triggering to the Cytoplasmic Juxtamembrane Regions of CD3zetazeta. Immunity, 43(2), 227–239. https://doi.org/10.1016/j.immuni.2015.06.018

Lewis, R. S. (2001). Calcium Signaling Mechanisms in T Lymphocytes. Annual Review of Immunology, 19(1), 497–521. https://doi.org/10.1146/annurev.immunol.19.1.497

Lin, J. J. Y., Low-Nam, S. T., Alfieri, K. N., McAffee, D. B., Fay, N. C., & Groves, J. T. (2019). Mapping the stochastic sequence of individual ligand-receptor binding events to cellular activation: T cells act on the rare events. Sci Signal, 12(564). https://doi.org/10.1126/scisignal.aat8715

Lo, W. L., Shah, N. H., Ahsan, N., Horkova, V., Stepanek, O., Salomon, A. R., Kuriyan, J., & Weiss, A. (2018). Lck promotes Zap70-dependent LAT phosphorylation by bridging Zap70 to LAT. Nat Immunol, 19(7), 733–741. https://doi.org/10.1038/s41590-018-0131-1

Lo, W. L., Shah, N. H., Rubin, S. A., Zhang, W., Horkova, V., Fallahee, I. R., Stepanek, O., Zon, L. I., Kuriyan, J., & Weiss, A. (2019). Slow phosphorylation of a tyrosine residue in LAT optimizes T cell ligand discrimination. Nat Immunol, 20(11), 1481–1493. https://doi.org/10.1038/s41590-019-0502-2

Lo, W. L., & Weiss, A. (2021). Adapting T Cell Receptor Ligand Discrimination Capability via LAT. Front Immunol, 12, 673196. https://doi.org/10.3389/fimmu.2021.673196

Mariuzza, R. A., Agnihotri, P., & Orban, J. (2020). The structural basis of T-cell receptor (TCR) activation: An enduring enigma. J Biol Chem, 295(4), 914–925. https://doi.org/10.1074/jbc.REV119.009411

Marunaka, Y. (2015). Roles of interstitial fluid pH in diabetes mellitus: Glycolysis and mitochondrial function. World J Diabetes, 6(1), 125–135. https://doi.org/10.4239/wjd.v6.i1.125

McAffee, D. B., O’Dair, M. K., Lin, J. J., Low-Nam, S. T., Wilhelm, K. B., Kim, S., Morita, S., & Groves, J. T. (2021). Discrete LAT condensates encode antigen information from single pMHC:TCR binding events. bioRxiv, 2021.2012.2016.472676. https://doi.org/10.1101/2021.12.16.472676

Michaux, A., Mauen, S., Breman, E., Dheur, M. S., Twyffels, L., Saerens, L., Jacques-Hespel, C., Gauthy, E., Agaugue, S., Gilham, D. E., & Sotiropoulou, P. A. (2022). Clinical Grade Manufacture of CYAD-101, a NKG2D-based, First in Class, Non-Gene-edited Allogeneic CAR T-Cell Therapy. J Immunother, 45(3), 150–161. https://doi.org/10.1097/CJI.0000000000000413

Mossman, K. D., Campi, G., Groves, J. T., & Dustin, M. L. (2005). Altered TCR signaling from geometrically repatterned immunological synapses. Science, 310(5751), 1191–1193. https://doi.org/10.1126/science.1119238

Nguyen, V. P., Alves, D. S., Scott, H. L., Davis, F. L., & Barrera, F. N. (2015). A Novel Soluble Peptide with pH-Responsive Membrane Insertion. Biochemistry, 54(43), 6567–6575. https://doi.org/10.1021/acs.biochem.5b00856

Nye, J. A., & Groves, J. T. (2008). Kinetic control of histidine-tagged protein surface density on supported lipid bilayers. Langmuir, 24(8), 4145–4149. https://doi.org/10.1021/la703788h

O’Donoghue, G. P., Pielak, R. M., Smoligovets, A. A., Lin, J. J., & Groves, J. T. (2013). Direct single molecule measurement of TCR triggering by agonist pMHC in living primary T cells. Elife, 2, e00778. https://doi.org/10.7554/eLife.00778

Pielak, R. M., O’Donoghue, G. P., Lin, J. J., Alfieri, K. N., Fay, N. C., Low-Nam, S. T., & Groves, J. T. (2017). Early T cell receptor signals globally modulate ligand:receptor affinities during antigen discrimination. Proc Natl Acad Sci U S A, 114(46), 12190–12195. https://doi.org/10.1073/pnas.1613140114

Prakaash, D., Cook, G. P., Acuto, O., & Kalli, A. C. (2021). Multi-scale simulations of the T cell receptor reveal its lipid interactions, dynamics and the arrangement of its cytoplasmic region. PLoS Comput Biol, 17(7), e1009232. https://doi.org/10.1371/journal.pcbi.1009232

Reinherz, E. L. (2019). The structure of a T-cell mechanosensor. Nature, 573(7775), 502–504. https://doi.org/10.1038/d41586-019-02646-w

Schamel, W. W., Alarcon, B., & Minguet, S. (2019). The TCR is an allosterically regulated macromolecular machinery changing its conformation while working. Immunol Rev, 291(1), 8–25. https://doi.org/10.1111/imr.12788

Scott, H. L., Heberle, F. A., Katsaras, J., & Barrera, F. N. (2019). Phosphatidylserine Asymmetry Promotes the Membrane Insertion of a Transmembrane Helix. Biophys J, 116(8), 1495–1506. https://doi.org/10.1016/j.bpj.2019.03.003

Scott, H. L., Westerfield, J. M., & Barrera, F. N. (2017). Determination of the Membrane Translocation pK of the pH-Low Insertion Peptide. Biophys J, 113(4), 869–879. https://doi.org/10.1016/j.bpj.2017.06.065

Smith, A. W., Smoligovets, A. A., & Groves, J. T. (2011). Patterned two-photon photoactivation illuminates spatial reorganization in live cells. J Phys Chem A, 115(16), 3867–3875. https://doi.org/10.1021/jp108295s

Tinevez, J. Y., Perry, N., Schindelin, J., Hoopes, G. M., Reynolds, G. D., Laplantine, E., Bednarek, S. Y., Shorte, S. L., & Eliceiri, K. W. (2017). TrackMate: An open and extensible platform for single-particle tracking. Methods, 115, 80–90. https://doi.org/10.1016/j.ymeth.2016.09.016

Trebak, M., & Kinet, J. P. (2019). Calcium signalling in T cells. Nat Rev Immunol, 19(3), 154–169. https://doi.org/10.1038/s41577-018-0110-7

Westerfield, J. M., Sahoo, A. R., Alves, D. S., Grau, B., Cameron, A., Maxwell, M., Schuster, J. A., Souza, P. C. T., Mingarro, I., Buck, M., & Barrera, F. N. (2021). Conformational Clamping by a Membrane Ligand Activates the EphA2 Receptor. J Mol Biol, 433(18), 167144. https://doi.org/10.1016/j.jmb.2021.167144

Wolpert, E. Z., Petersson, M., Chambers, B. J., Sandberg, J. K., Kiessling, R., Ljunggren, H. G., & Karre, K. (1997). Generation of CD8+ T cells specific for transporter associated with antigen processing deficient cells. Proc Natl Acad Sci U S A, 94(21), 11496–11501. https://doi.org/10.1073/pnas.94.21.11496

Wong, J., Obst, R., Correia-Neves, M., Losyev, G., Mathis, D., & Benoist, C. (2007). Adaptation of TCR repertoires to self-peptides in regulatory and nonregulatory CD4+ T cells. J Immunol, 178(11), 7032–7041. https://doi.org/10.4049/jimmunol.178.11.7032

Xu, C., Gagnon, E., Call, M. E., Schnell, J. R., Schwieters, C. D., Carman, C. V., Chou, J. J., & Wucherpfennig, K. W. (2008). Regulation of T cell receptor activation by dynamic membrane binding of the CD3epsilon cytoplasmic tyrosine-based motif. Cell, 135(4), 702–713. https://doi.org/10.1016/j.cell.2008.09.044

Yu, Y., Fay, N. C., Smoligovets, A. A., Wu, H. J., & Groves, J. T. (2012). Myosin IIA modulates T cell receptor transport and CasL phosphorylation during early immunological synapse formation. PLoS One, 7(2), e30704. https://doi.org/10.1371/journal.pone.0030704

Zhang, H., Cordoba, S. P., Dushek, O., & van der Merwe, P. A. (2011). Basic residues in the T-cell receptor zeta cytoplasmic domain mediate membrane association and modulate signaling. Proc Natl Acad Sci U S A, 108(48), 19323–19328. https://doi.org/10.1073/pnas.1108052108

