## Supplementary material for "Allosteric Inhibition of the T Cell Receptor by a Designed Membrane Ligand": Full supplementary materials

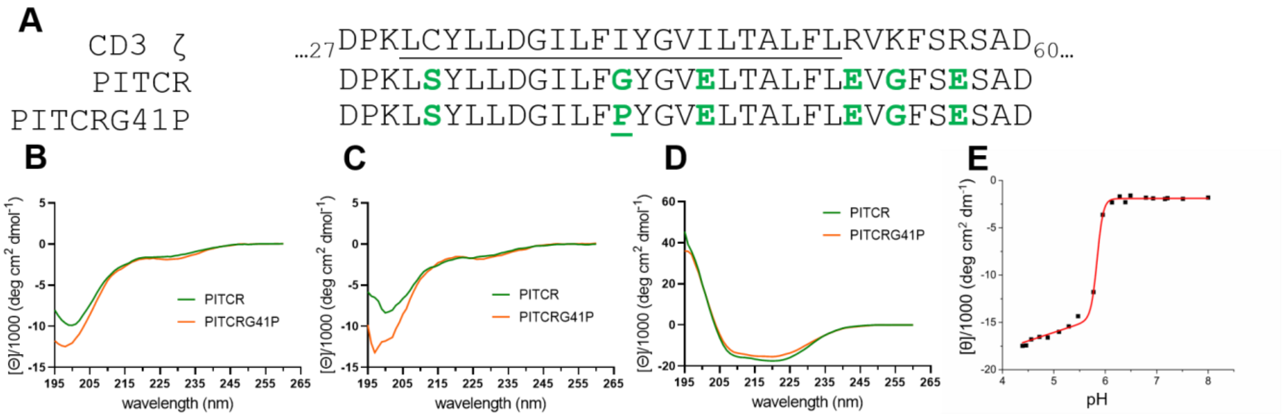


**Figure 1 - Figure supplementary 1. (A)** *Top*, the partial amino acid sequence from human CD3ζ comprising a small segment of the extracellular domain, the transmembrane domain (underlined) and a small portion of intracellular domain. *Middle*, the amino acid sequence of PITCR peptide. *Bottom*, the amino acid sequence of PITCRG41P. Introduced residues are highlighted in green. Residue numbers are labeled in human CD3ζ sequence. Circular dichroism spectra for PITCR and PITCRG41P in different conditions: 10 mM sodium phosphate buffer at pH 7.4 **(B)**, and in the presence of vesicles of 16:0-18:1 POPS/POPC (1/9) at pH 7.4 **(C)** and at pH 5.0 **(D)**. Each spectrum is the mean of three independent experiments. **(E)** Representative circular dichroism pH titration curve of PITCR in the presence of 16:0-18:1 POPS/POPC (1/9) liposomes. Data are calculated by the difference of molar ellipticity ([θ]) between 222 and 260 nm. Data are representative of two independent experiments. $pK_{CD}$ is 5.87 ± 0.03 (mean ± S.D.).

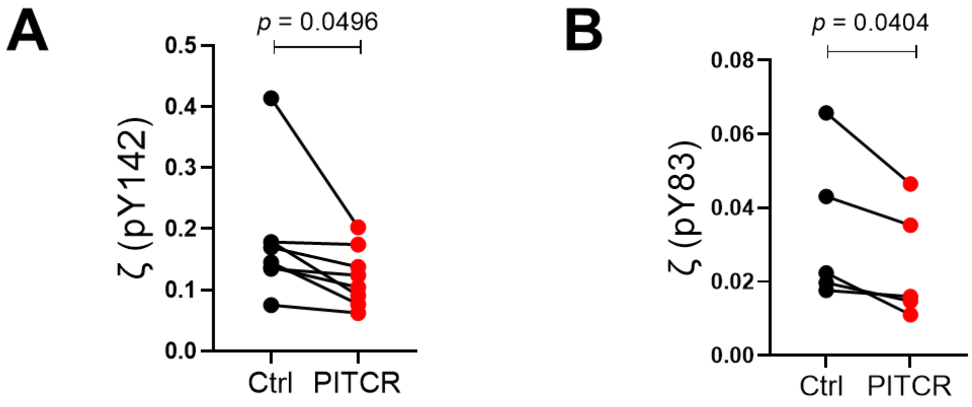


**Figure 1 - Figure supplementary 2. Quantification of phosphorylation of ζ (pY142)**. (A) and ζ (pY83) (B) after OKT3 stimulation. Band intensities were normalized to ζ (total). Each dot pair represents one independent experiment. *p* values were calculated using a two-tailed paired *t*-test.

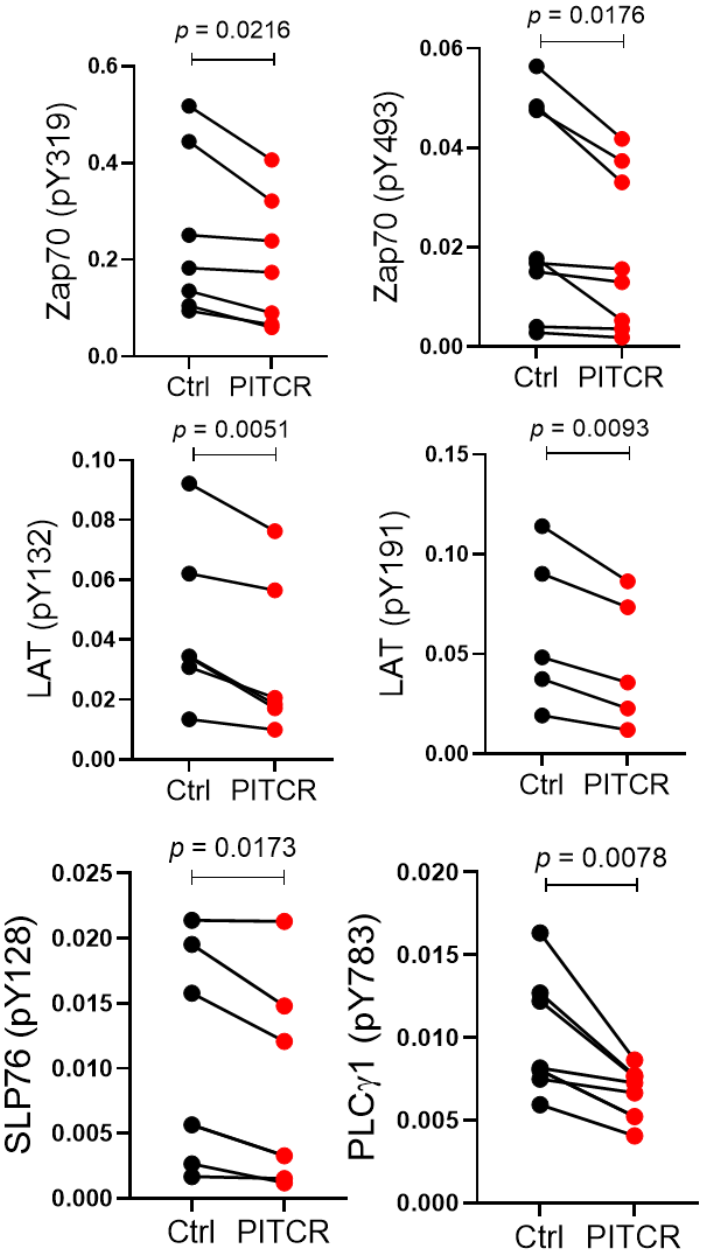


**Figure 2 - Figure supplementary 1. Quantification of Zap70 (pY319), Zap70 (pY493), LAT (pY132), LAT (pY191), SLP76 (pY128), and PLCγ1 (pY783) in response to OKT3 stimulation.** Zap70 (pY319) and Zap 70 (pY493) were normalized with Zap70 (total). LAT (pY132) and LAT (pY191) were normalized with LAT (total). SLP76 (pY128) and PLCγ1 (pY783) were normalized with β-actin. Each dot pair represents one independent experiment. All *p* values except PLCγ1 (pY783) were calculated using a two-tailed paired *t* test. *p* value for PLCγ1 (pY783) was calculated using a two-tailed Wilcoxon matched-pairs signed-rank test because the *p* value for the F test to compare variance was 0.0494.

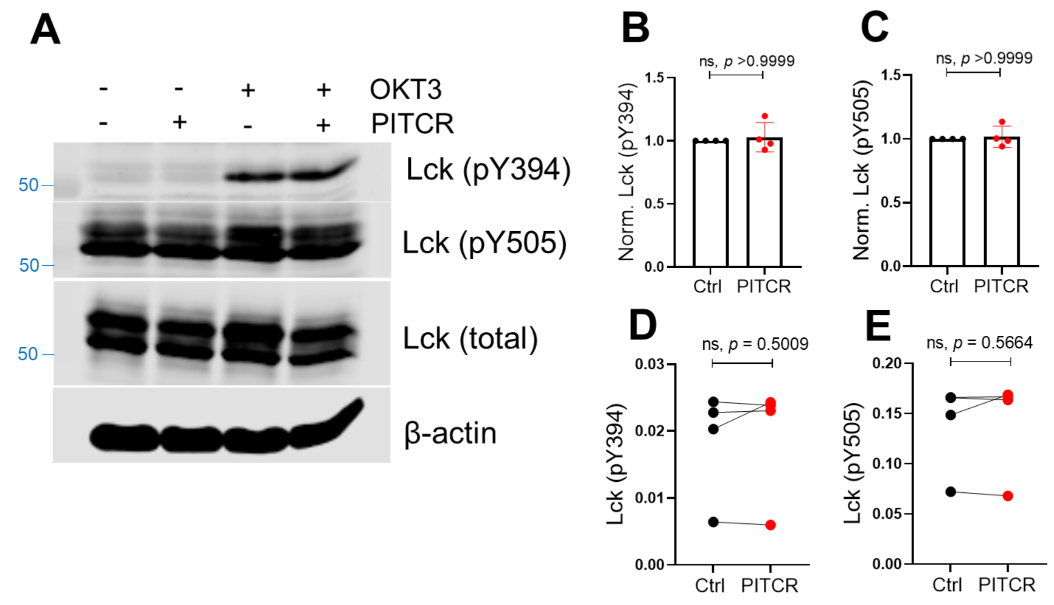


**Figure 2 - Figure supplementary 2. PITCR does not reduce phosphorylation of Lck after OKT3 activation.** (A) Immunoblot analysis of Lck (pY394 and pY505), Lck (total) and the housekeeping protein β-actin. Data are representative of four independent experiments. (B) and (C) show normalized quantifications of phosphorylation of tyrosine at the positions of 394 and 505 of Lck after TCR activation. Data in the presence of PITCR was normalized to data in the absence of PITCR in response to OKT3 activation. Error bars are the SD. *p* values were calculated using a two-tailed Mann-Whitney test. (D) and (E) show quantification of phosphorylation of Lck at pY394 and pY505 after the OKT3 stimulation. Band intensities were normalized to Lck (total). Each dot pair represents one independent experiment. *p* value was calculated using a two-tailed paired *t*-test.

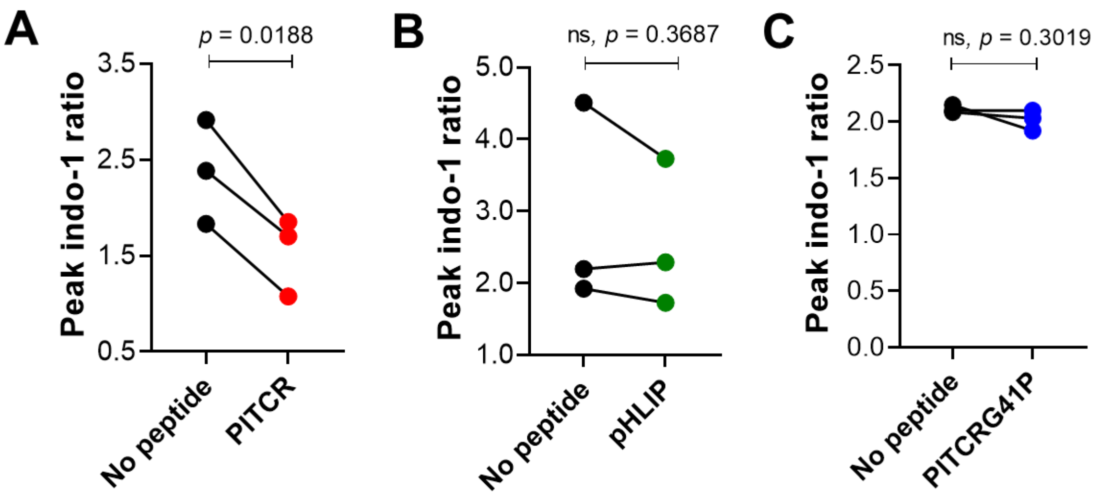


**Figure 3 - Figure supplementary 1. PITCR reduces intracellular calcium responses for PITCR (A), but not for pHLIP (B) or PITCRG41P (C**). Quantification of the magnitude of Indo-1 ratio between OKT3 peak and baseline. Each dot pair represents one independent experiment. Each independent experiment (n= 3) includes at least four technical replicates. *p* values were calculated using two-tailed paired Student's *t* test.

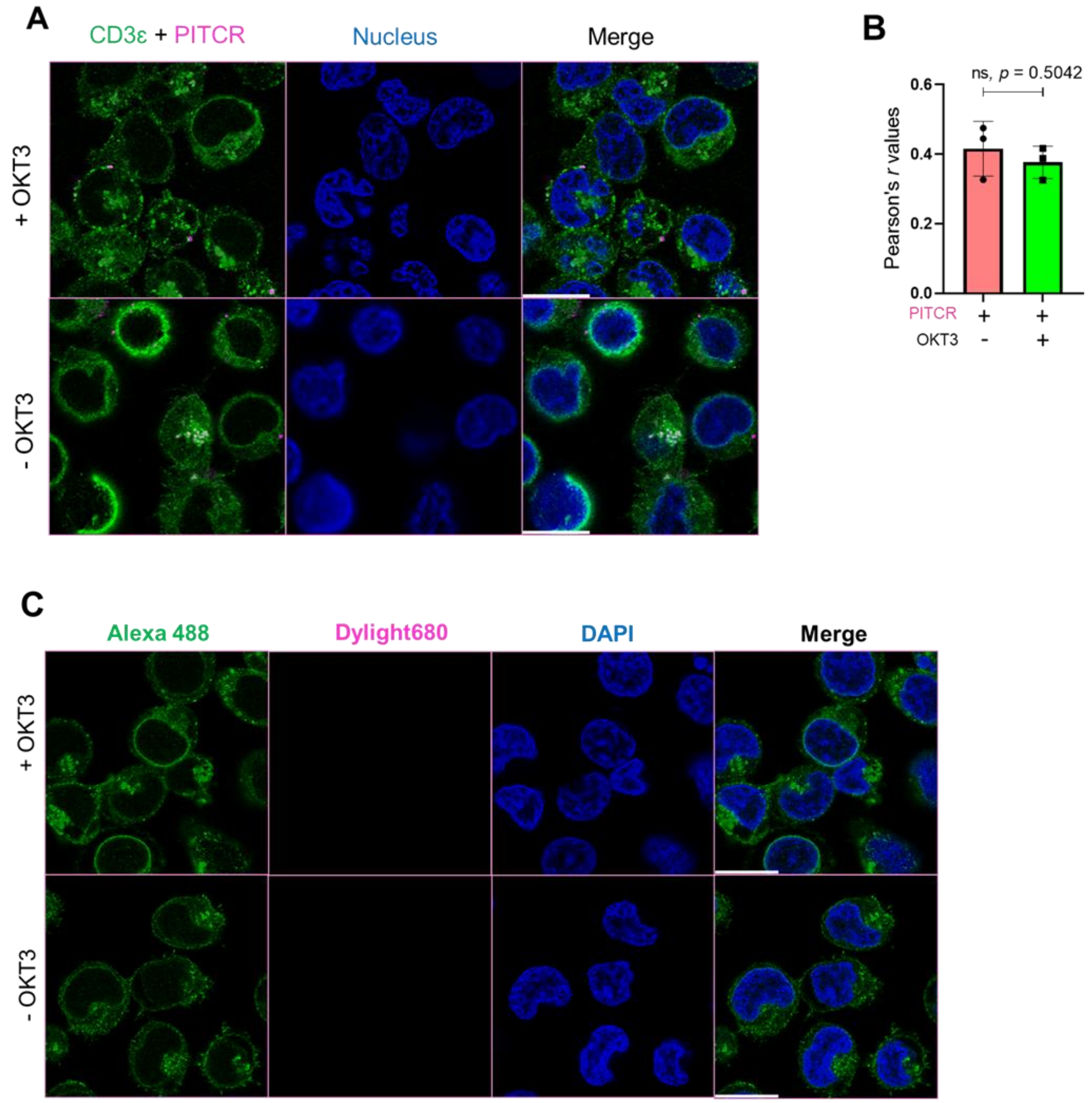


**Figure 5 Figure supplementary 1. PITCR colocalizes with TCR.** (A) PITCR and CD3ε co-localization images and nuclear staining (DAPI) are shown. Confocal images are representative of three independent experiments. Scale bars = 10 μm. (B) Quantification of PITCR and CD3ε co-localization calculating Pearson's *r* value. Each dot of panel B denotes one technical replicate from three independent experiments. N = 19-21. Error bars indicate SD. *p* value was calculated using a two tailed unpaired *t*-test. (C) Jurkat cells were treated with control followed by an anti-CD3ε immunofluorescent staining. Confocal images are representative of three independent experiments. Scale bars = 10 μm.

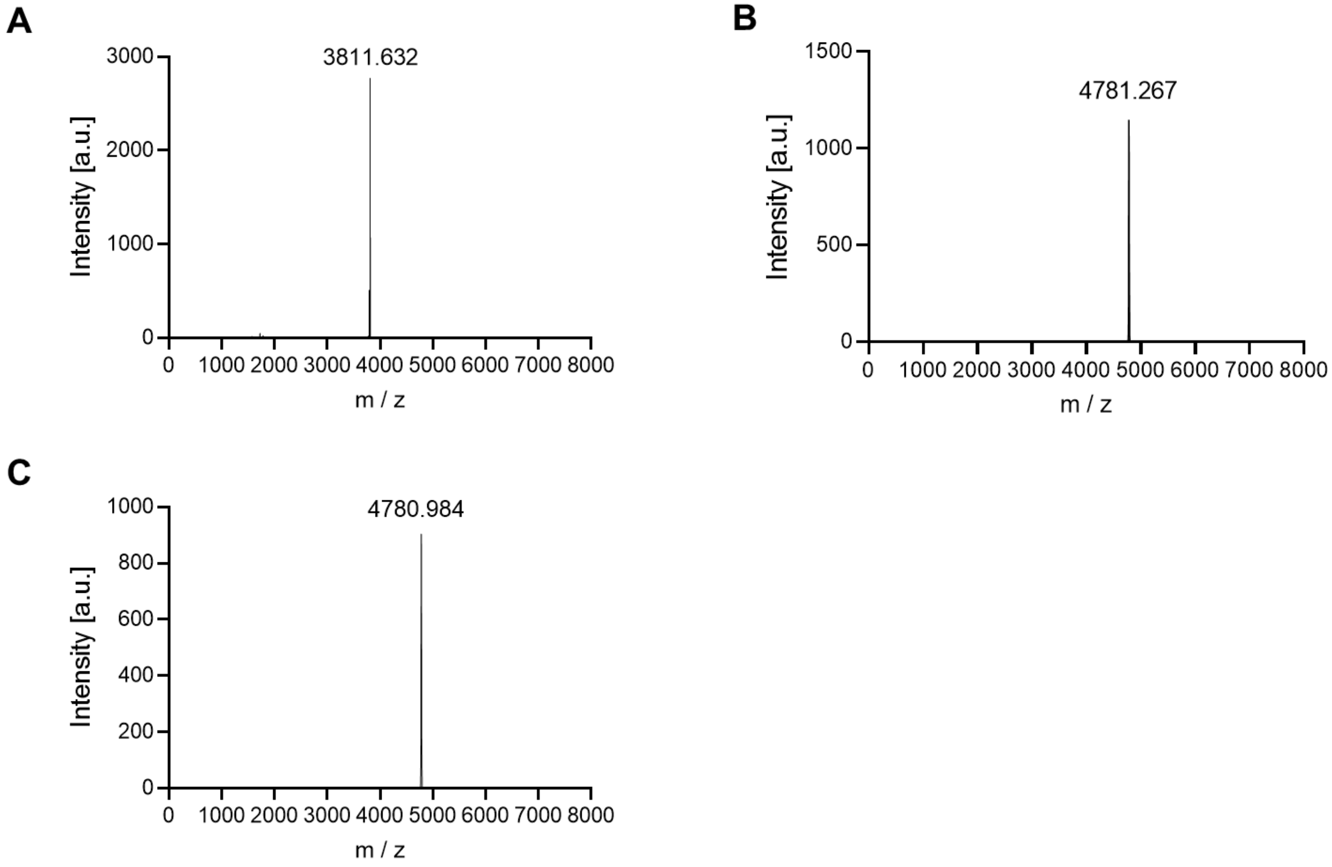


**Figure 5 - Figure supplementary 2. MALDI-TOF spectra of NEC-PITCR.** (A), PITCR conjugated with dylight680 (B), and PITCR conjugated with AZ555 (C). The theoretical MW of NEC-PITCR is 3812.31. The MW of dylight 680 and AZ555 are 972 Da and 969.12 Da, respectively.

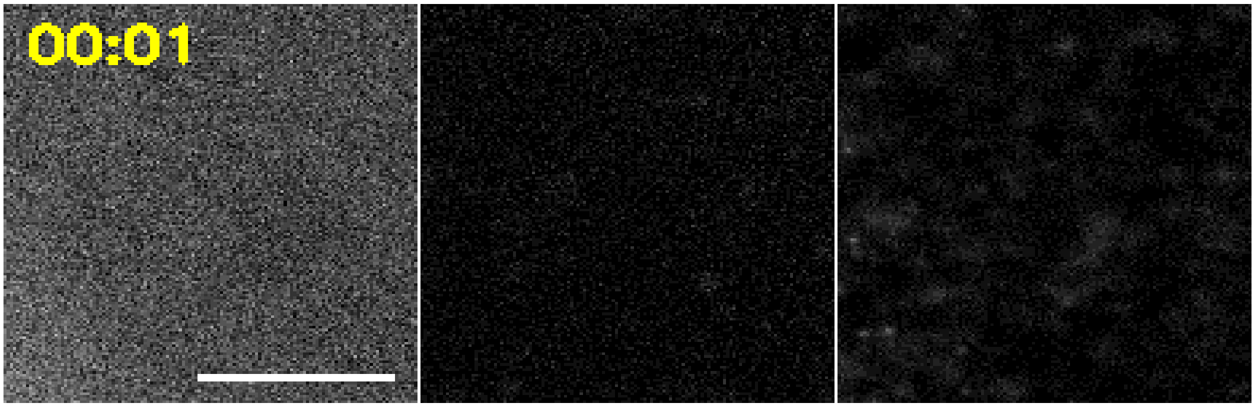


**Video 1: Real-time imaging of PITCR and pMHC in a T cell adhering to supported bilayer.** Left: RICM, center: PITCR555, right: pMHC. Cell footprint is shown as cyan line. Scale bar: 10 µm.

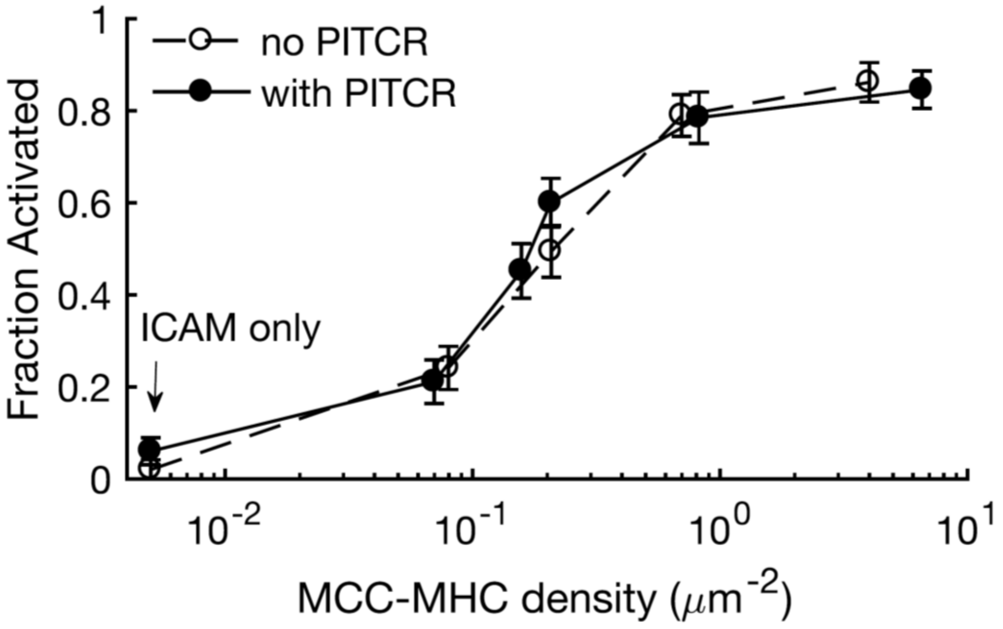


**Figure 6 - Figure supplementary 1. The NFAT dose-response curve of primary murine T cells is unaffected by PITCR.** AND-TCR primary murine CD4+ T cells expressing NFAT-mCherry reporter protein were stimulated by supported lipid bilayers functionalized with varied density of pMHC (MCC peptide 100% labeled with Atto-647N) and ICAM-1 (~20 molecules/µm2). Cells were defined as activated if the NFAT-mCherry signal in the nucleus was greater than the signal in the cytoplasm, as determined by epifluorescence imaging. The fraction of activated cells is indistinguishable between cells treated with or without PITCR at all pMHC densities. Error bars denote SEM.

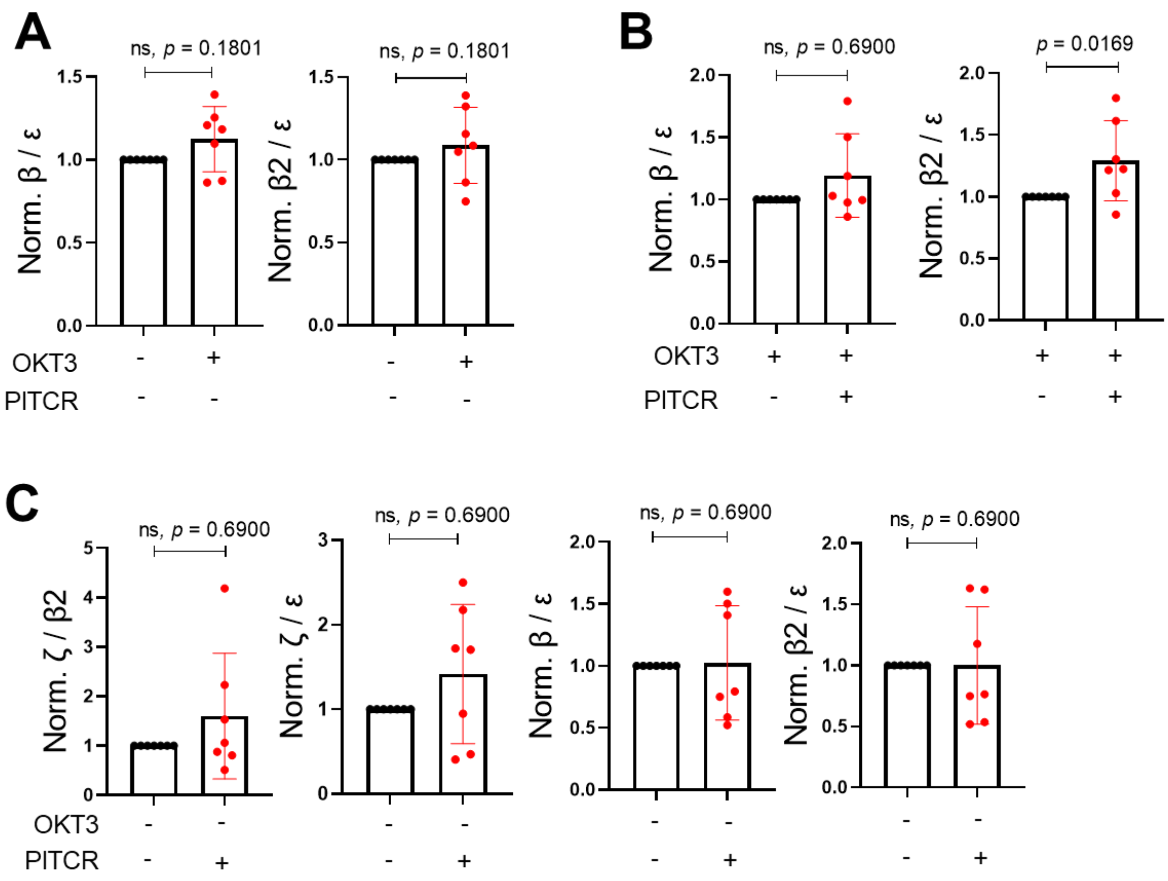


**Figure 8 Figure supplementary 1.** Quantification of DDM immunoprecipitation results for the following conditions: (A) OKT3 stimulation, (B) OKT3 stimulation in the presence or absence of PITCR and (C) PITCR incubation without stimulation. Error bars are SD. *p* values were calculated with a two-tailed Mann-Whitney test.
